# SOX2 utilizes FOXA1 as a heteromeric transcriptional partner to drive proliferation in therapy-resistant prostate cancer

**DOI:** 10.1101/2025.07.18.664790

**Authors:** John T. Phoenix, Audris Budreika, Devin A. Schmeck, Raymond J. Kostlan, Marina G. Ferrari, Kristen S. Young, Charles S. Rogers, Carleen D. Deegan, Marcus W. Bienko, Hannah E. Bergom, Ella Boytim, Ryan M. Brown, Julia A. Walewicz, Shreya K. Bhagi, Leigh Ellis, Emmanuel S. Antonarakis, Justin M. Drake, Pushpinder S. Bawa, Jordan E. Vellky, Anthony Williams, Natalie M. Rezine, Jonathan P. Rennhack, Sean W. Fanning, Justin H. Hwang, Russell Z. Szmulewitz, Donald J. Vander Griend, Steven Kregel

## Abstract

Treatment options and diagnostic outlook for men with advanced, therapy resistant prostate cancer (PCa) are extremely poor; this is primarily due to the common lack of durable response to androgen receptor (AR) targeted therapies and phenotypic transdifferentiation into a particularly lethal subtype known as neuroendocrine prostate cancer (NEPC). In this study, we mechanistically determine that SOX2 (a transcription factor originally repressed by AR) physically binds and acts in a concerted manner with FOXA1 (a key AR pioneering cofactor) to regulate a subset of genes which promote cell cycle progression, and lineage plasticity in AR-refractory prostate cancers. Our findings assert the SOX2/FOXA1 interaction as an important mediator of resistance to AR-targeted therapy and a driver of NEPC and lineage plasticity; their coordinated action and downstream signaling offers a potential novel therapeutic opportunity in late-stage PCa.

## Introduction

One in 44 men die of prostate cancer (PCa). It is the second leading cause of cancer-related deaths in men^1^. For nearly nine decades, the androgen signaling axis has remained the foremost therapeutic target in PCa.^2^. In the normal prostate, a hormone-responsive transcription factor known as the androgen receptor (AR) regulates differentiation and lineage commitment^3,4^, primarily through an interaction with its requisite pioneering cofactor Forkhead box A1 (FOXA1)^5^. During carcinogenesis, AR switches toward oncogenic activity that can be successfully ablated with treatments such as androgen depletion (castration) or direct AR-targeted therapies^3,6,7^. However, in up to 20% of men, these treatments will fail within five years^8^, which ultimately leads to progressive metastatic disease and death^9–11^. Prolonged treatment with AR-targeted therapies can select for aggressive, lineage-plastic pathologic subtypes, the best characterized of which is acutely lethal neuroendocrine prostate cancer (NEPC). NEPC lacks canonical AR activity due to low/absent AR protein expression, but maintains *FOXA1* expression, which is essential for its survival^12^.

NEPC tumors are often defined by “de-differentiation” from their original prostate lineage. They express markers of neuroendocrine cells derived from a common prostate stem cell^13^ and express progenitor cell markers, many of which were initially repressed by AR during differentiation^14^ and are thus turned back on in the NEPC state. Previously, we identified *SOX2* [SRY (sex determining region Y)-box 2] as one such gene whose expression is directly repressed by AR^15^ in AR-driven prostate adenocarcinoma cells. SOX2 is a minor groove DNA-binding transcription factor, canonically known for its essential role in maintaining the survival and pluripotency of undifferentiated human embryonic stem cells (hESCs)^16,17^. However, SOX2 has an emerging role as an epigenetic reprogramming factor and oncogene^18–22^. In hESCs, SOX2 regulates the expression of roughly 1500 genes, many of which are co-regulated with OCT4 and/or Nanog^17^. In adult tissues, *SOX2* expression is found in cells within the basal-epithelial cell layer of normal glandular epithelia^23^ as well as in prostate tumors^15,23,24^. *SOX2* expression in prostate cancer is induced upon AR-inhibition, which maintains and promotes the proliferation and survival of prostate cancer cells^15,25,26^. Its expression in prostate cancer is correlated with increased Gleason grade, increased metastasis, decreased time to biochemical recurrence, and resistance to AR-targeted therapy^15,25,27,28^.

Like many SOX-family transcription factors, SOX2 must heterodimerize with another transcription factor to bind DNA; its heterodimer partner is tissue-specific and determines target gene specificity^29^. The heterodimeric partner within hESCs is OCT3/4^30^; however, SOX2’s requisite binding partner(s) in PCa remains largely unknown. Our recent studies revealed that SOX2 has a unique cistrome in castration-resistant PCa (CRPC) cells compared to hESCs^25^. Based on this finding, we hypothesized that SOX2 binds as a heterodimer to another transcription factor in order to promote oncogenic activity in late-stage prostate cancer. Through motif scanning of the SOX2 cistrome, we prioritized FOXA1; a Forkhead box (FOX) family protein that serves as a pioneering transcription factor for many nuclear hormone receptors, including AR^5^. FOXA1 is an oncogene in PCa that is essential for AR activity in adenocarcinoma ^5,31,32^ and NEPC survival^12^. In this study we employed biochemical approaches to detect protein-protein interactions, assayed PCa viability after SOX2 knockdown, and assessed genomic occupancy of AR, FOXA1, and SOX2 in adenocarcinoma and NEPC contexts. Overall, our experiments aimed to test the hypothesis that SOX2 utilizes FOXA1 as its heterodimeric partner in prostate cancer, driving a pathologic, oncogenic switch from FOXA1’s normal behavior of binding AR.

Our results demonstrate that SOX2 and FOXA1 engage in direct physical interaction in both AR-expressing but castration-resistant adenocarcinoma and NEPC cell lines via co-immunoprecipitation (Co-IP), proximity ligation assay (PLA), *in silico* structural modeling, and a split nano-luciferase assay (Nano-BiT). *SOX2* knockdown invokes a proliferation defect which significantly decreases survival of both adenocarcinoma and NEPC cells *in vitro*. To our knowledge, this is the first evidence of a direct SOX2 dependency within a representative NEPC system. In both adenocarcinoma and NEPC models, we find ∼90% overlap between SOX2 and FOXA1 binding sites. These co-bound regions are also AR-regulated in adenocarcinoma cells, and are functionally enriched for cell cycle progression genes, many of which have been shown to promote lineage plasticity and neuroendocrine differentiation. When looking at NEPC patient datasets, we show a decoupling of AR and FOXA1-regulated gene networks and an establishment of SOX2 and FOXA1-regulated gene networks. Furthermore, we show that SOX2 seldom shares the same binding sites in cancers compared to its canonical environments of pluripotent and prostate basal epithelial cells (PrECs). This suggests a unique, cancer-specific DNA binding profile of SOX2 alongside FOXA1 in malignant disease. Prostate cancer’s bypass of current therapeutic interventions is of great clinical interest; here, we mechanistically show the oncogenic SOX2-FOXA1 interaction and its unique cistrome can facilitate cell survival, resistance to AR-targeted therapies, and lineage plastic forms of the disease.

## Results

### SOX2 knockdown decreases survival of SOX2-positive adenocarcinoma and neuroendocrine prostate cancer cells

First, we compared the expression of SOX2 in a panel of immortalized non-tumorigenic patient-derived prostate epithelial cells (PrEC), AR-expressing adenocarcinoma, and AR-negative prostate cancer cells, including the NEPC cell line NCI-H660 (H660) (**Figure 1A**). As we identified previously, *SOX2* expression is limited to CWR-R1 and CWR-22Rv1 (22Rv1)^15^ among the tested adenocarcinoma cell lines and is highest in the NEPC cell line NCI-H660 (**Figure 1A**). Additionally, another cell line model of NEPC, LASCPC-01^33^, had comparable levels of SOX2 (**Figure 1B**). FOXA1 was expressed in all cell lines, regardless of subtype or *AR* expression (**Figure 1A and 1B**).

**Figure 1.**
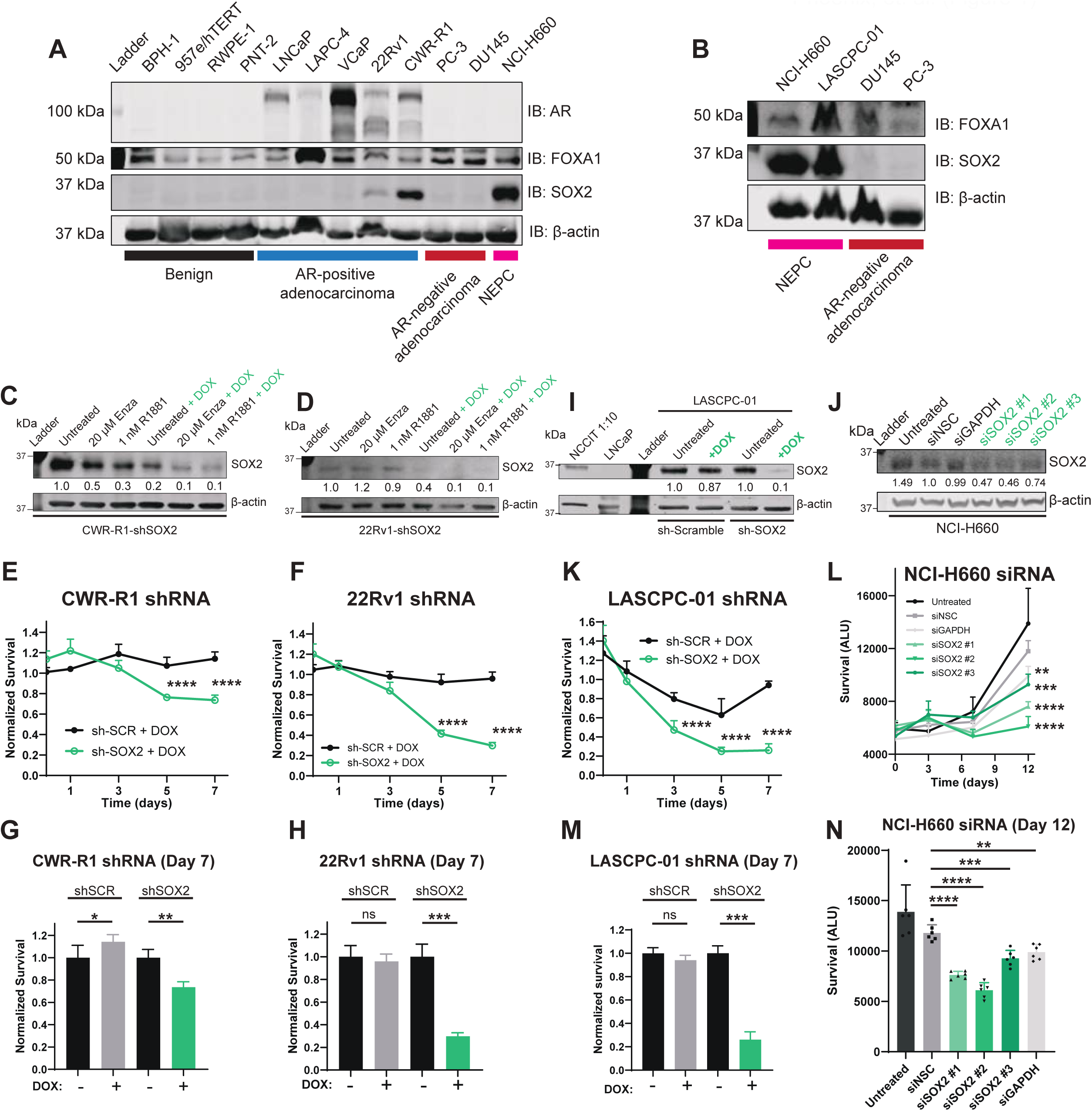
SOX2 knockdown decreases survival of SOX2-positive adenocarcinoma and neuroendocrine prostate cancer cells. **A)** Western blot panels of both non tumorigenic and malignant prostate cells for AR, FOXA1, and SOX2. Blots for β-actin were used as an internal loading control. **B)** Western blot panels of AR-negative PCa cell lines for AR, FOXA1, SOX2. Blots for β-actin were used as an internal loading control. **C-D)** Western blots show successful SOX2 shRNA knockdown at 48h in CWR-R1 and 22Rv1 adenocarcinoma cells. Protein bands were quantified using Empiria Studio. **E-F)** Normalized survival plots of CWR-R1 and 22Rv1 cells after doxycycline induction of shRNA targeting *SOX2* or a non-specific scrambled shRNA (sh-SCR). Data points represent the mean normalized survival ± SD; each point is normalized to its own untreated control (without DOX) at each time interval. **G-H)** Bar graphs represent normalized survival of both CWR-R1 and 22Rv1 cells 7 days after dox-induction of shRNA. **I-J)** Western blot show successful SOX2 knockdown at 48h via shRNA in LASCPC-01 and siRNA in NCI-H660 NEPC cells. Protein bands were quantified using Empiria Studio. **K)** Normalized survival plots of LASCPC-01 NEPC cells after doxycycline induction of shRNA targeting *SOX2* or a non-specific scrambled shRNA (sh-SCR). Data points represent the mean normalized survival ± SD; each point is normalized to its own untreated control (without DOX) at each time interval. **L)** Survival plot of NCI-H660 NEPC cells treated with siRNAs targeting SOX2. Data points represent mean luminescence +/- SD. **M)** Bar graph represents normalized survival of LASCPC-01 NEPC cells 7 days after dox-induction of shRNA. **N)** Bar graph represents raw survival of NCI-H660 NEPC cells 12 days after siRNA transfection. Data points represent mean luminescence +/- SD.

We next sought to identify the functional consequence of SOX2 depletion across PCa contexts. To do so, we utilized doxycycline-inducible short-hairpin RNA (shRNA) against *SOX2* in CWR-R1 and 22Rv1 prostate adenocarcinoma cells. In both CWR-R1 and 22Rv1, we found that SOX2 knockdown was successful regardless of AR signaling modulation **(Figure 1C-D and S1A-B)**. We observed a significant decrease in cell viability in the SOX2 knockdown group compared to the group treated with a non-specific scrambled control shRNA (shSCR) **(Figure 1E and 1F)**, with their largest deficiency in normalized cell survival occurring seven days after the initial shRNA induction **(Figure 1G and 1H)**. Next, to assay the effect of SOX2 knockdown in the AR-negative NEPC context, we employed a similar dox-inducible shRNA system against *SOX2* in LASCPC-01, alongside an alternative approach of *SOX2*-targeting siRNAs in H660 given the technical difficulty of lentiviral transduction in these cells. In both LASCPC-01 and H660, we found that SOX2 protein was successfully knocked down after treatment **(Figure 1I and 1J)**, resulting in a significant decrease in longitudinal cell survival in the SOX2 knockdown groups compared to their respective non-specific controls **(Figure 1K and 1L)**. The impact of SOX2 knockdown on NEPC cell survival was most pronounced in LASCPC-01 seven days post-induction **(Figure 1M)** and 12 days post-transfection in H660 **(Figure 1N)**. When observing the non-normalized growth curves of CWR-R1, 22Rv1, and LASCPC-01, SOX2 knockdown imparts a clear defect in cell proliferation across time points (**Figures S1C-E**) suggesting a mechanism in which SOX2 is required for both adenocarcinoma and NEPC cells to proliferate, independent of AR status.

### SOX2 directly interacts with FOXA1 in PCa cells in a manner competitive with AR, and this is corroborated with expression data in PCa patient samples

Our prior ChIP-seq experiments on SOX2 in CWR-R1 adenocarcinoma cells show that SOX2 does not display a similar global cistrome as it does in WA-01 hESC cells^25^, implying that in the absence of OCT3/4, SOX2 utilizes another binding partner to facilitate unique chromatin interactions and function in PCa. We used this data to identify candidate heterodimeric transcription factor partners within 20 nucleotide base pairs of a SOX2 peak through motif scanning analysis. We identified several FOX motifs enriched near SOX2 peaks, including the canonical FOXA1 motif (**Figure 2A**). On average, the FOXA1 motif is 10 base pairs away from the canonical SOX2 motif (**Figure 2A**), which implies dimerization along the same face of the DNA double helix, and is the exact periodicity across the nucleosome exhibited in canonical SOX2-OCT3/4 interactions in hESCs^34,35^.

**Figure 2.**
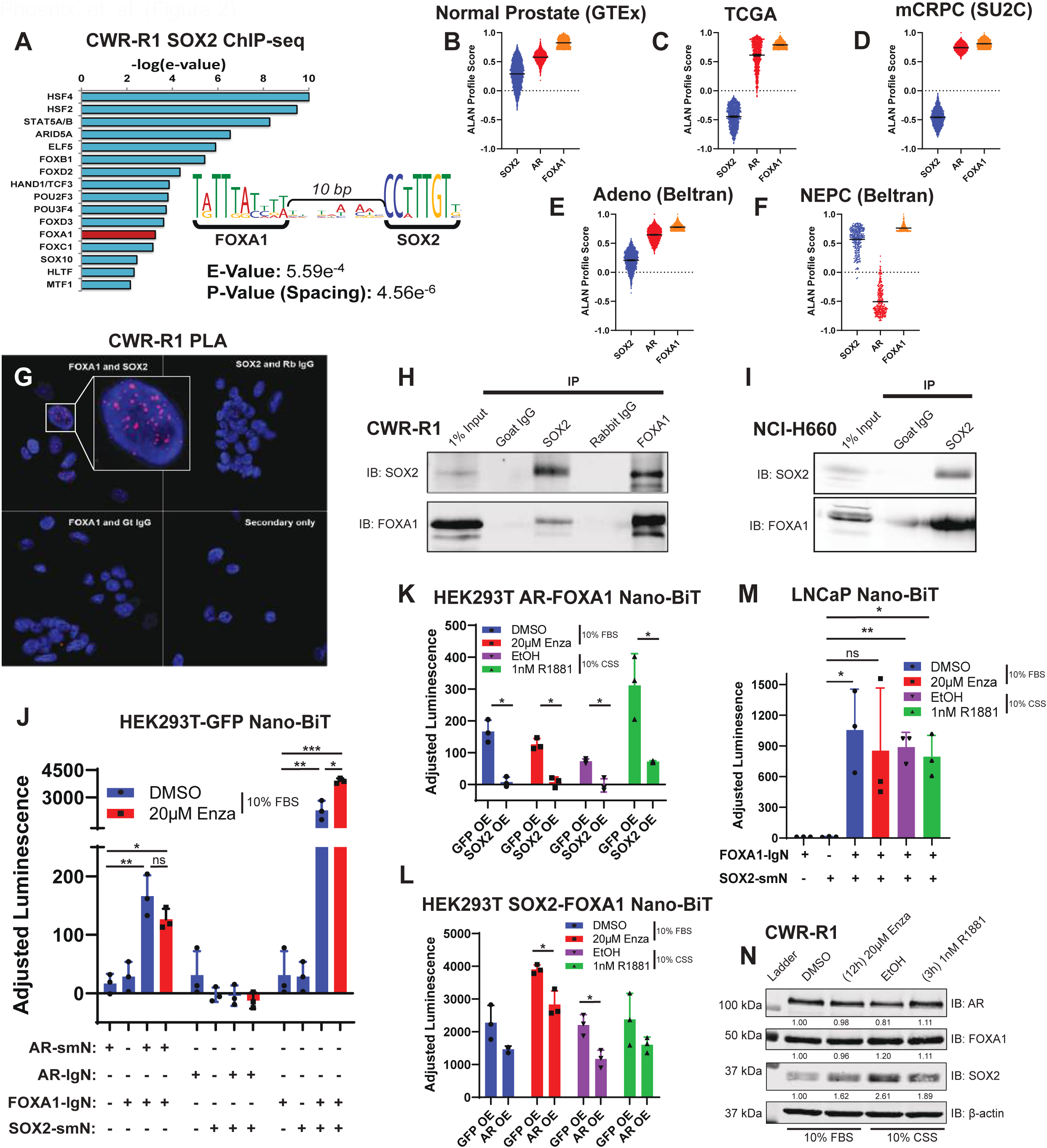
SOX2 directly interacts with FOXA1 in PCa cells competitively with AR, as is corroborated with gene expression data in PCa patient samples. **A)** All candidate TFs within 20bp of SOX2 peak in CWR-R1 cells. The e-value is the lowest p-value of any spacing of the secondary motif times the number of secondary motifs; it estimates the expected number of random secondary motifs that would have the observed minimum p-value or less. FOXA1 identified in red. The FOXA1-SOX2 motif enriched by ChIP-seq shows a potential direct interaction on the chromatin. **B-F)** Gene network signature analyses in PCa patient tumors using the Algorithm for Linking Activity Networks (ALAN) model. Datasets from publicly available sources are depicted across contexts of prostate cancer (GTEX, TCGA, SU2C, and Adeno and NEPC). Gene network signature for FOXA1 (orange) is visualized compared to the gene networks of AR (blue) and SOX2 (red) in each dataset where the median ALAN profile score for each gene network is noted with a black line. Overall network analysis is represented as positive for a median ALAN profile score above 0 and negative for a median ALAN profile score below 0. **G)** Proximity ligation assay (PLA) measures co-localization (red) of SOX2 and FOXA1 CWR-R1 castration-resistant prostate adenocarcinoma cells. **H)** Reciprocal co-immunoprecipitation of SOX2 and FOXA1 in CWR-R1 cells. **I)** Co-immunoprecipitation of FOXA1 with SOX2 in the NEPC cell line NCI-H660. **J)** Split nano-luciferase complementation reporter assay (Nano-BiT) demonstrating a specific interaction between SOX2 and FOXA1 in HEK293T cells. Tagged AR and FOXA1 were shown to interact as a positive control, and co-transfection of tagged AR and SOX2 were a negative control. Data points represent mean luminescence +/- SD. **K)** Lentiviral *SOX2* overexpression (OE) ablates the AR-FOXA1 Nano-BiT interaction in HEK293T cells. Data points represent mean luminescence +/- SD. **L)** Lentiviral *AR* overexpression (OE) ablates the SOX2- FOXA1 Nano-BiT interaction in HEK293T cells, specifically in ARSI conditions. Data points represent mean luminescence +/- SD. **M)** Nano-BiT in prostate adenocarcinoma LNCaP cells demonstrates strong SOX2-FOXA1 interaction. No significant changes were observed across different AR-signaling contexts. Data points represent mean luminescence +/- SD. **N)** Western blots of CWR-R1 protein lysate from cells that underwent AR-pathway modulation. Protein bands were quantified using Empiria Studio.

Given FOXA1’s maintained expression and essential oncogenic role in prostate cancer^5,31,32^, we looked to patient transcriptome datasets to see how *FOXA1* expression and gene behavior compares to both SOX2 and AR by utilizing the Algorithm for Linking Activity Networks (ALAN)^36^ model. Bulk normal prostate tissue sequencing (GTEX dataset)^37^ suggests that AR, FOXA1, and SOX2 gene network activity is correlated (**Figure 2B and S2H**). However, given that these proteins are differentially expressed in the varying cell types within normal prostate tissue, SOX2 in the basal cells, and AR and FOXA1 in the luminal cells^5,15^, bulk assessments of expression alone would make it difficult to assess the SOX2-FOXA1 interaction. In localized PCa that is largely AR-driven (TCGA dataset)^38^, AR and FOXA1 gene network signatures are more closely aligned with each other and less so with SOX2 (**Figure 2C and S2I**), an effect which is even more pronounced in the metastatic CRPC dataset, enriched for amplified or mutated AR (SU2C dataset)^39^ (**Figure 2D and S2J**). However, in high-grade prostate adenocarcinoma patients enriched for tumors that have adjacent neuroendocrine features [Adeno (Beltran) dataset]^40^, the ALAN scores of AR, SOX2 and FOXA1 begin to correlate (**Figure 2E and S2K**). Strikingly, in NEPC tumors [NEPC (Beltran) dataset]^41^, we observe a sharp decoupling of the AR gene network signature from FOXA1, and the establishment of a strong SOX2 and FOXA1 gene network relationship (**Figure 2F and S2L**). This suggests that SOX2 may promote lineage plasticity, mediating both AR-independent growth and the NEPC phenotype through an interaction with FOXA1.

To investigate if SOX2 and FOXA1 physically bind each other, we utilized a variety of biochemical approaches to assay protein-protein interactions. We performed both proximity ligation assay (PLA) and co-immunoprecipitation (Co-IP) in our cell line model of advanced CRPC (CWR-R1) and observed SOX2 and FOXA1 physically near one another (**Figure 2G and 2H**). We then confirmed SOX2/FOXA1 interaction in the cell line model of NEPC (H660) via Co-IP (**Figure 2I**). As a confirmation of direct SOX2/FOXA1 interaction, we established a split nano-luciferase complementation reporter protein-protein interaction assay (Nano-BiT) in HEK293T cells that were engineered to overexpress AR, SOX2, or GFP as a control (**Figure S2A-C**)^42^. Through optimization of the split-luciferase tag locations at either the N- or C-termini, SOX2 and FOXA1 interact strongly with N-terminally tagged constructs (**Figure 2J**), and the values of SOX2-FOXA1 interaction far exceed our positive control AR and FOXA1 interaction, and we detect no direct interaction with SOX2 and AR (**Figure 2J**). These data strongly suggest direct binding of SOX2 and FOXA1. Given the degree to which the SOX2-FOXA1 protein interaction BiT signal exceeded the AR-FOXA1 signal, we then tested the hypothesis that SOX2 could outcompete AR binding to FOXA1. We observe that lentiviral SOX2 overexpression completely abolishes AR-FOXA1 Nano-BiT signal in HEK293T cells under all conditions aside from short-term androgen stimulation (1nM R1881, 3h treatment) in which SOX2 overexpression still decreases AR-FOXA1 interaction by ∼75% (**Figure 2K**). Whereas SOX2 overexpression leads to near-complete elimination of AR-FOXA1 interactions experimentally, AR overexpression has minimal effect on the SOX2-FOXA1 interaction, with a statistical reduction only seen with AR ligand deprivation (CSS) or antagonism (20 μM enzalutamide) (**Figure 2L**). In LNCaP adenocarcinoma PCa cells, which endogenously express high levels of AR, we find that SOX2 and FOXA1 interact via Nano-BiT regardless of hormone manipulation when SOX2 is exogenously expressed (**Figure 2M**). These data suggest that the presence of SOX2, even in AR-expressing PCa, could hijack FOXA1 and alter gene expression.

Since SOX2 associates with FOXA1 in prostate cancer cells, we next hypothesized that SOX2 might have preferential binding to FOXA1 displaying clinically relevant pathogenic mutations, specifically the well-characterized R219C/S form of FOXA1^31,43–45^. We show that SOX2 still binds to FOXA1 harboring R219 mutations, but we observe no changes in affinity compared to wild-type as assayed through Nano-BiT in AR-overexpressing HEK293T cells (**Figure S2D**). Finally, we modulated AR activity in CWR-R1 adenocarcinoma cells, which are known to co-express all three proteins, and assayed the effects on AR, FOXA1, and SOX2 protein levels. As we had previously identified, *SOX2* expression was induced by AR inhibition (20 μM enzalutamide or hormone depletion via CSS)^15^ (**Figure 2N**). However, heightened SOX2 levels decrease after only 3 hours of androgen stimulation (**Figure 2N**), suggesting that aside from AR repressing SOX2 transcriptionally, it may also directly outcompete SOX2 from a transcriptional complex with FOXA1. In terms of PLA signal, we see increased SOX2/FOXA1 signal in CWR-R1 cells treated with enzalutamide and see decreased SOX2/FOXA1 signal in cells treated with 1nM R1881 (**Figure S2E-G**), which mirrors this effect (**Figure 2N)**. This perhaps occurs through enhanced AR interaction with other cofactors, resulting in decreased SOX2 stability in cells. Taken together, these data show a strong, direct SOX2/FOXA1 interaction even when AR is present and active, and regardless of FOXA1 mutation status. As such, cistrome analyses are warranted to understand the behavioral relationship between these proteins on a chromatin level.

### SOX2 co-occupies sites on DNA with both FOXA1 and AR in CRPC adenocarcinoma cells, specifically near genes that promote cell proliferation

Next, we investigated SOX2, FOXA1, and AR chromatin binding through ChIP-seq, first in CWR-R1 adenocarcinoma cells. We performed ChIP-seq under our four different AR-modulation conditions: normal growth conditions (10% FBS), 24-hour treatment with 20μM enzalutamide, 24-hour hormone starvation (CSS), and short-term AR activation (3-hour treatment with 1nM R1881). As expected, we identified AR (full site – Consensus Androgen Response element) and FOXA1 (half-site – ARE) motifs with AR immunoprecipitation (**Figure 3A**). We identified both FOX and SOX motifs enriched near FOXA1 binding sites, in addition to significant enrichment of motifs containing the core SOX2 binding motif [TTGT] (**Figure 3A**). Lastly, near SOX2 binding peaks we find significant enrichment of both AR and FOXA1 motifs **(Figure 3A)**. When we compared the binding of AR, FOXA1, and SOX2 to each other, we unsurprisingly find that about 2/3 of AR peaks overlap at least 1 base pair with FOXA1 peaks, which is typical for prostate cancer cells^31^ (**Figure 3B**). However, we found an even greater overlap between SOX2 peaks and FOXA1 peaks (83.9%), in addition to the vast majority of the SOX2 peaks (77.6%) being co-localized with AR as well (**Figure 3B**).

**Figure 3.**
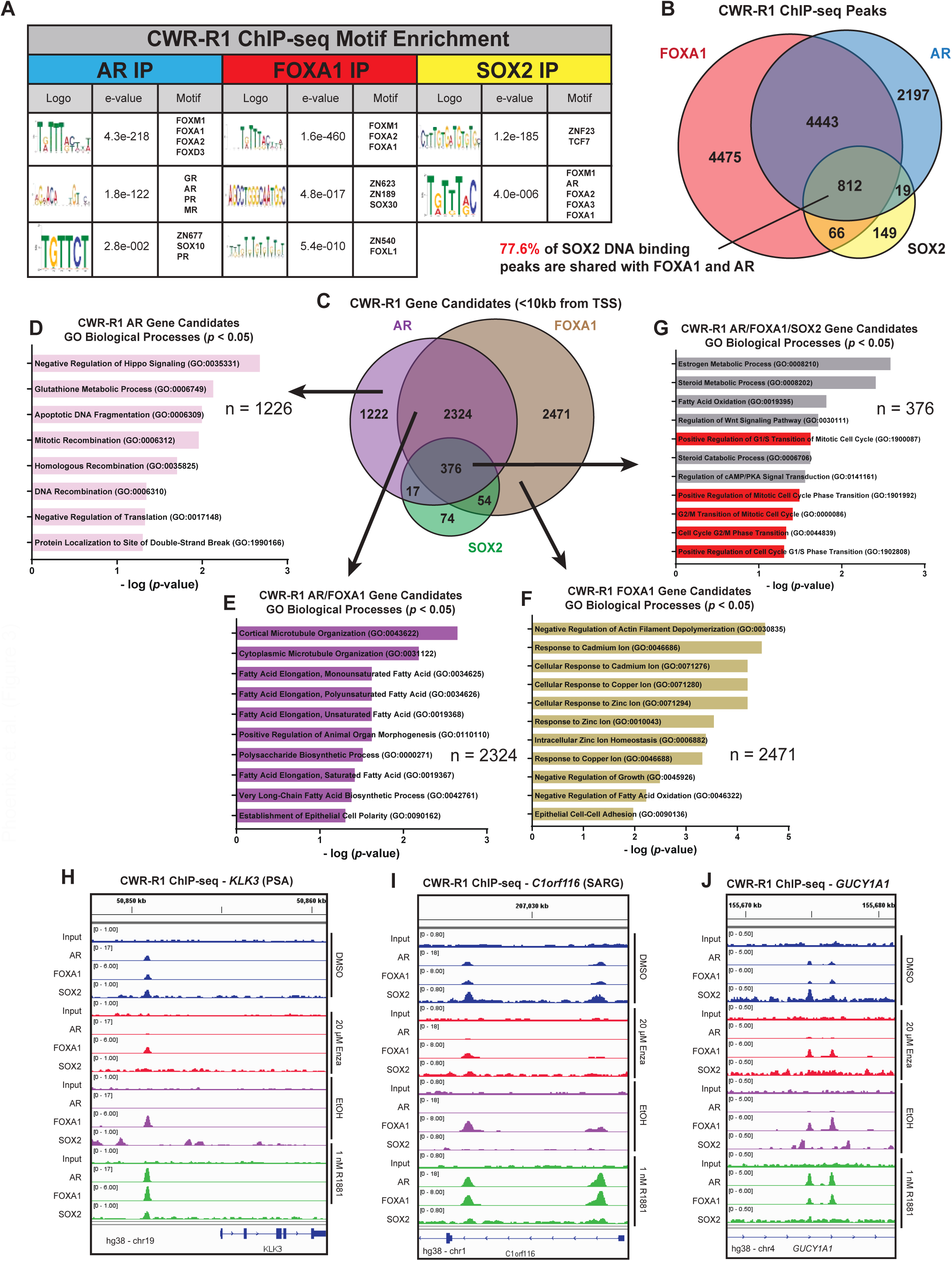
SOX2 co-occupies sites on DNA with both FOXA1 and AR in CRPC adenocarcinoma cells, specifically near genes that promote cell proliferation. **A)** Binding motif enrichment near AR, FOXA1, and SOX2 peaks in CWR-R1 cells. Both MEME and STREME motif discovery analyses were included. Full lists of motifs discovered can be found in Supplementary Table 1. **B) C)** Venn diagram of AR, FOXA1, and SOX2 ChIP-seq peaks in CWR- R1 cells grown in whole media. Shared regions represent called peaks with >= 1bp of overlap. **D)** Venn diagram of AR, FOXA1, and SOX2 potential target genes. CWR-R1 ChIP-seq peaks were analyzed using Cistrome-GO to identify genes that had AR, FOXA1, and/or SOX2 transcription factor peaks less than 10kb from a TSS and an adjusted regulatory potential (RP) score of >0.01. **E-H)** Gene Ontology (GO) pathway analyses for the unique and shared AR, FOXA1, and SOX2 candidate target genes in CWR-R1. GO enrichment was performed using Enrichr. **I-K)** Track plots of AR, FOXA1, and SOX2 ChIP-seq peaks in CWR-R1 undergoing AR pathway modulation. Peaks visualized using IGV and mapped to the human hg38 chromosome build.

To assess which genes these transcription factors might be regulating, we filtered these raw ChIP-seq peaks to select for peaks within 10kb of gene transcriptional start sites (TSS) using Cistrome-GO and observed a similar overlap profile between SOX2, FOXA1, and AR (**Figure 3C**). We then performed Gene Ontology (GO) analysis on the subset of genes near AR peaks alone (**Figure 3D**), genes near FOXA1 and AR overlapping peaks (**Figure 3E**), genes near FOXA1 peaks alone (**Figure 3F**) and on genes near SOX2, FOXA1, and AR overlapping peaks (**Figure 3G**). Strikingly, the subset of genes near SOX2, FOXA1, and AR co-bound regions (72.2% of the total SOX2 peaks) were functionally enriched for GO (Biological processes) terms involved in proliferation and cell cycle progression (**Figure 3G**); and these GO terms did not occur without SOX2 binding (**Figures 3D-F**) (see full list in **Figure S3**). Rather than proliferation, the genes enriched near AR peaks alone produced GO terms mostly involved in DNA repair processes (**Figure 3D**). AR genes co-bound with FOXA1 were enriched for GO terms involved in fatty acid synthesis and cytoskeletal rearrangement (**Figure 3E**), and genes enriched near FOXA1 peaks alone show potential regulation of metal ion transport (**Figure 3F**). Additionally, we identified SOX2 co-binding at known AR-regulated genes, such as PSA/*KLK3* (**Figure 3H**), SARG/*C1orf116* (**Figure 3I**), and *GUCY1A1* (**Figure 3J**), which we previously identified as co-regulated by AR and FOXA1 at selective androgen receptor response (sARE) sites^46^. These data suggest that SOX2 may bind to FOXA1 at AR co-regulated sites and could facilitate unique transcription of genes involved in cell proliferation.

### SOX2 and FOXA1 share a cistrome in NEPC and co-regulate key genes implicated in cell proliferation and lineage plasticity

Turning our attention to the cellular model of NEPC, which lacks AR entirely and has high endogenous levels of *SOX2* expression, we investigated SOX2 and FOXA1 genomic occupation in H660 cells via ChIP-seq. As was the case in adenocarcinoma, we identified an enrichment of FOXA1 binding motifs near SOX2 peaks, as well as enrichment of SOX motifs and the core [TTGT] nucleotides^25^ near FOXA1 peaks (**Figure 4A**). We also see an even more pronounced DNA binding overlap between SOX2 and FOXA1, with nearly 90% of SOX2 DNA binding peaks overlapping with FOXA1 (**Figure 4B**). As we did with CWR-R1 previously in **Figure 3**, we filtered the raw H660 ChIP-seq peaks to select for peaks within 10kb of gene transcriptional start sites (TSS) using Cistrome-GO and observed substantial overlap between SOX2 and FOXA1 candidate target genes (**Figure 4C**). Genes near FOXA1 peaks alone seem to be enriched for amino acid transport and differentiation processes (**Figure 4D**). SOX2 binding peaks alone, unsurprisingly, seem to map near genes involved in developmental processes (**Figure 4E**), and the overlapping SOX2/FOXA1 binding peaks are functionally enriched for many neuronal developmental and proliferation-associated GO terms, such as axon guidance (**Figure 4F**) (see full list in **Figure S4**). This finding supports what other groups have established to govern the NEPC phenotype^47–49^; a transcriptome profile that mirrors neuronal development.

**Figure 4.**
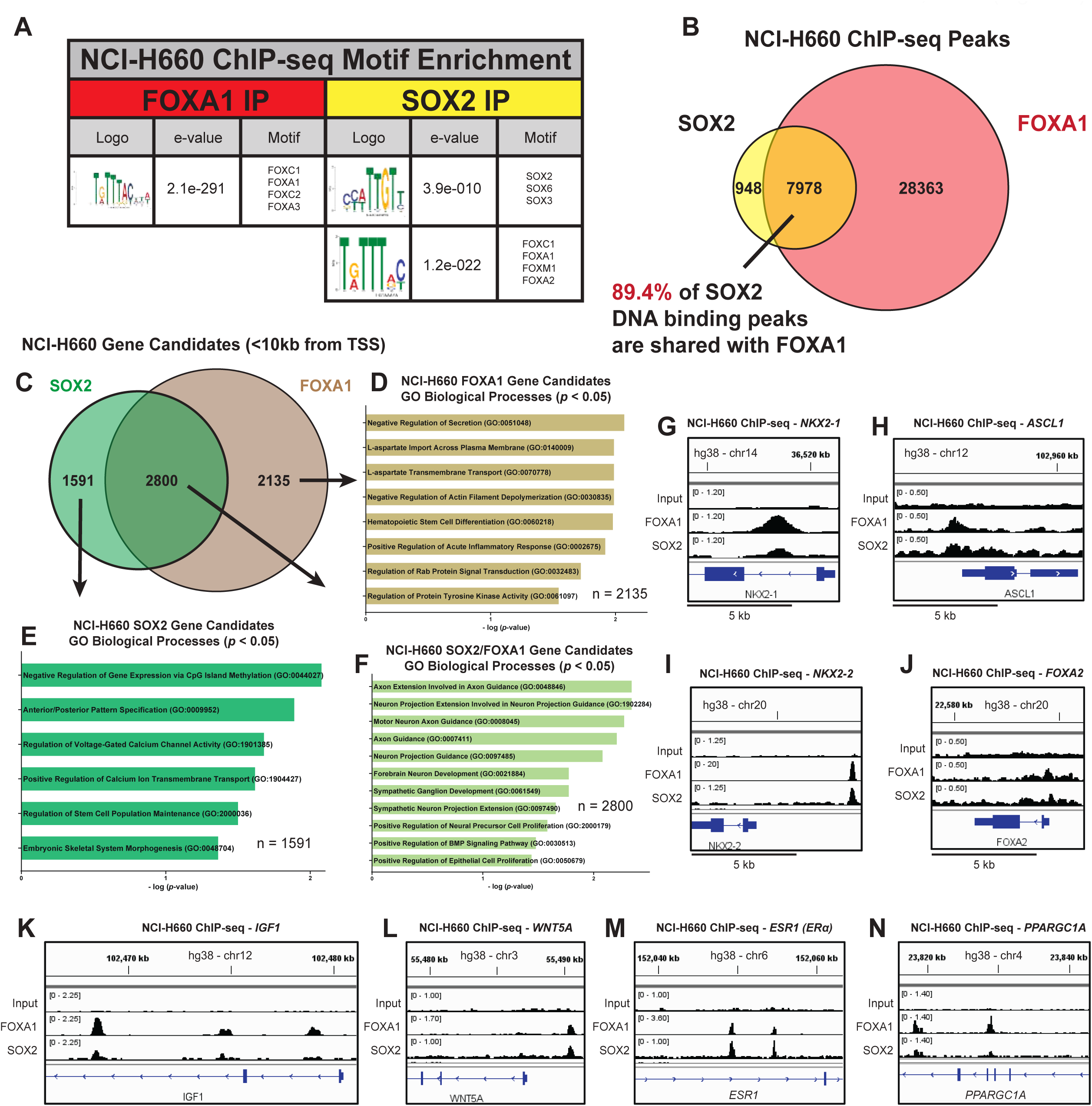
SOX2 and FOXA1 share a cistrome in NEPC and co-regulate key genes implicated in cell proliferation and lineage plasticity. **A)** Binding motif enrichment near FOXA1 and SOX2 peaks in NCI-H660 NEPC cells. Both MEME and STREME motif discovery analyses were included. Full lists of motifs discovered can be found in Supplementary Table 1. **B)** Venn diagram of SOX2 and FOXA1 ChIP-seq peaks in NCI-H660 cells grown in complete 5% FBS HITES media. Shared regions represent called peaks with >= 1bp of overlap. **C)** Venn diagram of SOX2 and FOXA1 potential target genes. NCI-H660 ChIP-seq peaks were analyzed using Cistrome-GO to identify genes that had AR, FOXA1, and/or SOX2 transcription factor peaks less than 10kb from a TSS and an adjusted regulatory potential (RP) score of >0.01. **D-F)** Gene Ontology (GO) pathway analyses for the unique and shared FOXA1 and SOX2 candidate target genes in NCI- H660. GO enrichment was performed using Enrichr. **G-N)** Track plots of SOX2 and FOXA1 ChIP- seq peaks near oncogenic/NEPC genes in NCI-H660. Peaks visualized using IGV and mapped to the human hg38 chromosome build.

Therefore, we then investigated our SOX2/FOXA1 co-occurring ChIP-seq peaks and identified many genes implicated in NEPC formation and lineage plasticity. SOX2 and FOXA1 co-bound sites were enriched near genes of important NEPC transcription factors, such as: *NKX2-1*^50^ (**Figure 4G**), *ASCL1*^51^ (**Figure 4H**), *NKX2-2*^47^ (**Figure 4I**), and *FOXA2*^52,53^ (**Figure 4J**). In addition to these targets, we also see SOX2 and FOXA1 co-localizing near genes such as *RET*, *PROX1*, and *TFAP4*, each of which are demonstrated drivers of PCa therapy resistance and the NEPC phenotype^54–56^ (**Figure S5A-C**). We confirm *FGF5* as a SOX2 target, which we previously found upregulated by SOX2 overexpression^15^ (**Figure S5D**), and identify co-binding at the important NEPC-related oncogene *IGF1*^57^ (**Figure 4K**). SOX2/FOXA1 co-bound peaks were enriched near *WNT5A*^58^ (**Figure 4L**) and its receptors, the receptor tyrosine kinase-like orphan receptors (*ROR2*, **Figure S5E** and *ROR1*, **Figure 6D**), whose signaling axis has been implicated in promoting NEPC^59^ and enzalutamide resistance in PCa^60^. Furthermore, SOX2 and FOXA1 are co-bound upstream of *SEMA3C*, whose expression is known to be driven by clinically relevant R219-mutated FOXA1 to promote CRPC (**Figure S5F**)^61^. Interestingly, SOX2/FOXA1 are also co-bound within the estrogen receptor alpha (ERα, *ESR1*) gene (**Figure 4M**), which is known to be highly upregulated in NEPC when compared to adenocarcinoma^57^. Lastly, SOX2 and FOXA1 bind together near the *PPARGC1A* gene, which has recently been established to drive small-cell neuroendocrine cancer progression toward an *ASCL1*-expressing subtype, a commonly observed and lethal phenomenon in NEPC^62^ (**Figure 4N**). Altogether, these data suggest that SOX2 and FOXA1 bind chromatin together to potentially mediate the expression of key genes that govern the NEPC phenotype and promote lineage plasticity in PCa. This suggests SOX2 and FOXA1 govern the NEPC phenotype, particularly via upstream regulation of ASCL1 and its associated genes

### SOX2 and FOXA1 reciprocally bind DNA sites and regulate proliferation in both adenocarcinoma and neuroendocrine prostate cancer cells

Given that SOX2 and FOXA1 co-bind with AR near genes that regulate proliferation in adenocarcinoma cells, and that in NEPC SOX2/FOXA1 co-bind near many genes that are important for the NEPC phenotype, we next sought to compare the findings of the CWR-R1 adenocarcinoma ChIP-seq to the H660 NEPC ChIP-seq. DNA binding heatmaps ( **Figure 5)** show reciprocal overlap between FOXA1 and SOX2 (**Figure 5A and 5B**) in CWR-R1, and vice versa in H660 (**Figure 5C and 5D**). We observe the most substantial binding overlap occurring by FOXA1 strongly occupying the top SOX2 peaks in both CWR-R1 and H660 (**Figure 5B and 5D**). When comparing the subset of shared SOX2/FOXA1 peaks within 10kb of gene transcriptional start sites (TSS) in either CWR-R1 or H660, we observed a conserved overlap of genes (130) that have SOX2/FOXA1 co-binding nearby in both CWR-R1 and H660. (**Figure 5E**). Functional GO enrichment of the NEPC-specific SOX2-FOXA1 co-bound candidate genes (**Figure 5F**) identifies many neuron-specific processes, as was the case in **Figure 4F**. Conversely, when analyzing the adenocarcinoma-specific SOX2-FOXA1 co-bound candidate genes (**Figure 5G**), we observe enrichment of many of the same androgen and steroid-regulation processes as were discovered in **Figure 3H**. However, the 130 genes co-bound by SOX2 and FOXA1 across cell lines show functional enrichment for genes involved in proliferation, cell-cycle progression, and the Wnt-signaling pathway (**Figure 5H**). Despite their distinct phenotypes, these data suggest irrespective of CRPC subtype (adenocarcinoma or NEPC), the SOX2-FOXA1 interaction promotes proliferation in PCa. This oncogenic interaction may be a central CRPC-enabling mechanism and useful target in both settings.

**Figure 5.**
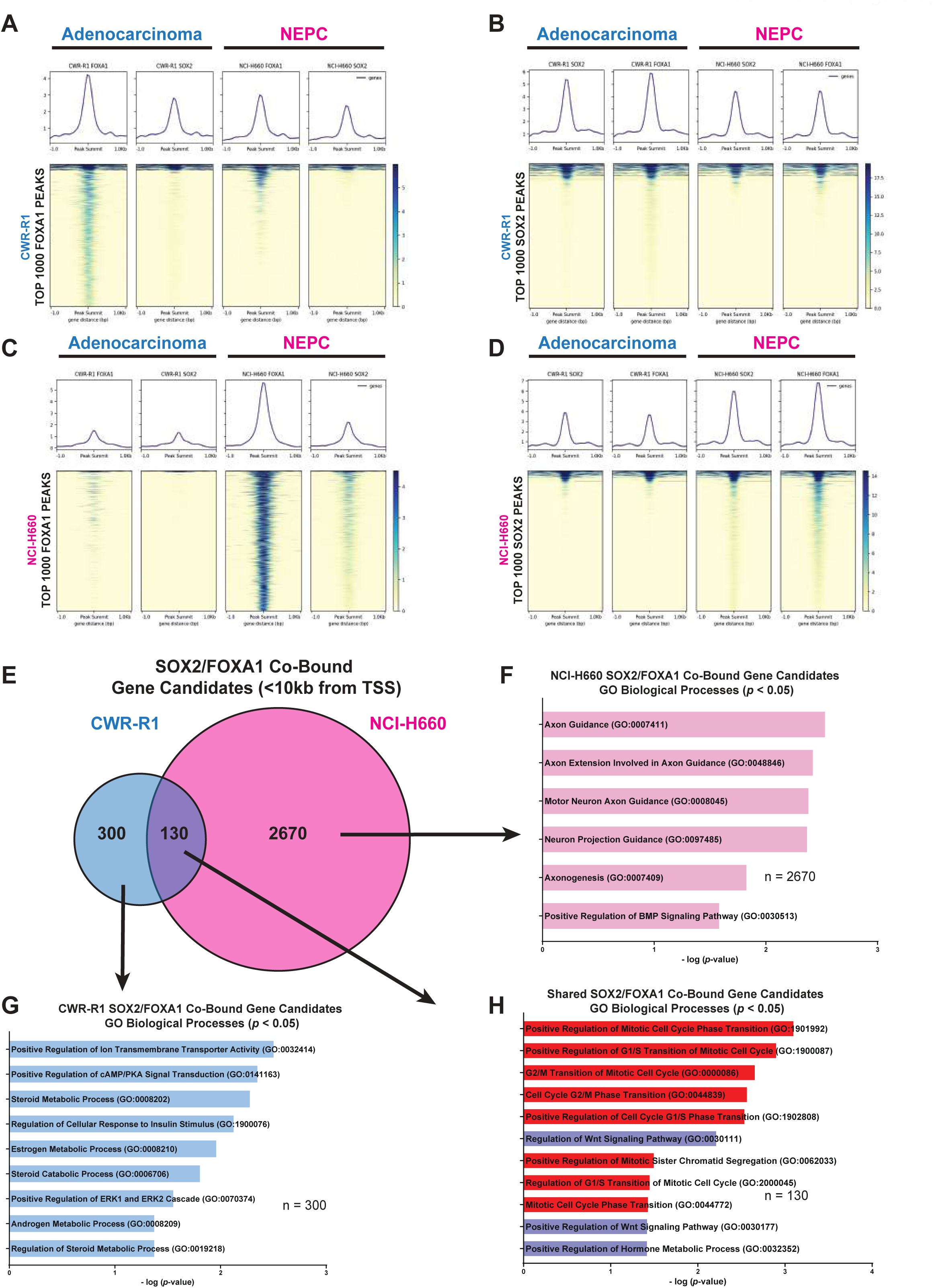
SOX2 and FOXA1 reciprocally bind DNA sites and regulate proliferation in both adenocarcinoma and NEPC contexts. **A-D**) Heatmaps of transcription factor binding in CWR- R1 and NCI-H660 cells. The vertical axis of each heatmap represents the top 1000 transcription factor binding sites by peak score in both adenocarcinoma and NEPC. **E)** Nested Venn diagram of the shared SOX2/FOXA1 candidate target genes (from Figures 3C and 4C) in both CWR-R1 and NCI-H660. We find 130 potentially SOX2/FOXA1 co-regulated genes across cell lines. **F-H**) Gene Ontology (GO) pathway analyses for the unique and shared CWR-R1 and NCI-H660 candidate SOX2/FOXA1 co-regulated genes. GO enrichment was performed using Enrichr.

**Figure 6.**
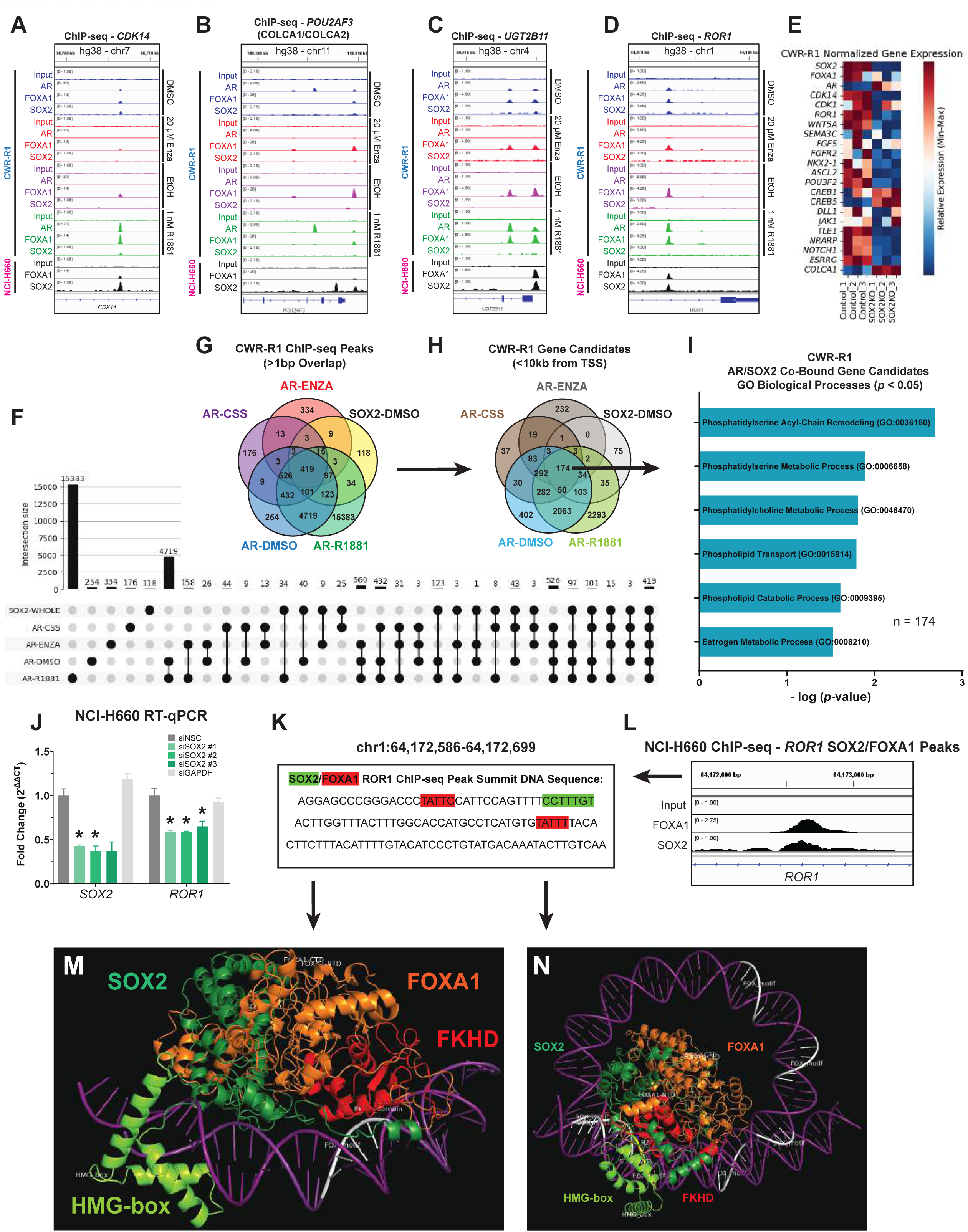
SOX2 co-occupies sites with AR that are unaffected by AR signaling modulation, suggesting stabilizing the complex on the chromatin to promote AR-signaling inhibition (ARSI) resistance and lineage plasticity. **A-D**) Track plots of AR, FOXA1, and SOX2 co-bound genes in CWR-R1 and NCI-H660. Peaks visualized using IGV and mapped to the human hg38 chromosome build. **E)** Heatmap of differential gene expression in Control vs. SOX2^KO^ CWR-R1 cells. Data represent transcripts per million (TPM) values of RNA-seq triplicates. **F)** UpSet plot of AR and SOX2 ChIP-seq peaks in CWR-R1 cells. Shared regions represent called peaks with >= 1bp of overlap. **G)** Venn Diagram of AR and SOX2 ChIP-seq peaks in CWR-R1 cells. Shared regions represent called peaks with >= 1bp of overlap. **H)** Venn diagram of AR and SOX2 potential target genes. CWR-R1 ChIP-seq peaks were analyzed using Cistrome-GO to identify genes that had AR and/or SOX2 binding peaks less than 10kb from a TSS and an adjusted regulatory potential (RP) score of >0.01. **I)** Gene Ontology (GO) pathway analyses for the invariant AR and SOX2 candidate target genes in CWR-R1. GO enrichment was performed using Enrichr. **J)** Relative *SOX2* and *ROR1* mRNA expression in NCI-H660 cells following *SOX2* siRNA knockdown normalized to β-actin. Data are represented as the fold change (2^-ΔΔCT^) ± SEM. **K)** The chosen sequence of the SOX2/FOXA1 co-bound region (peak summit) of the *ROR1* gene used for computational modeling of the SOX2/FOXA1 interaction. **L)** Track plot of SOX2/FOXA1 co-binding in the *ROR1* gene body. **M)** Computational modeling using the Chai-1 build of AlphaFold-3 to predict the structure of SOX2 and FOXA1 (UniProt IDs: P48431 and P55317, respectively) co-bound to DNA containing a single FOX motif within the *ROR1* gene. **N)** Structure prediction of SOX2 and FOXA1 co-bound to DNA containing both an HMG-box and FOX motif within the *ROR1* gene.

### SOX2 co-occupies sites with AR that are unaffected by AR signaling modulation, suggesting stabilizing the complex on the chromatin to promote AR-signaling inhibition (ARSI) resistance

Given the striking subset of genes at which SOX2 and FOXA1 co-localize in both adenocarcinoma and NEPC, we then set out to investigate some of these genes in further detail. Particularly, we focused on genes in CWR-R1 that are also co-bound by AR, to assess how SOX2 could potentially compensate for AR inhibition by utilizing FOXA1 as a partner. Many genes that AR binds in the adenocarcinoma setting continue to be bound by SOX2 in NEPC after *AR* expression is lost. This suggests maintenance of AR target gene expression by SOX2 and FOXA1 in the AR-negative PCa context. Some of the genes at which we identified SOX2 binding in both H660 and CWR-R1 are known to promote cell-cycle progression, such as *CDK14*^63^ (**Figure 6A**), as well as the *POU2AF3* locus (**Figure 6B**). The *POU2AF3* locus encodes the AR-responsive long non-coding RNA (lncRNA) COLCA1^64^ and the anti-sense transcript encodes COLCA2, a protein implicated in neuroendocrine differentiation^47^ and vital to the Tuft-cell lineage of small-cell neuroendocrine lung tumors^65^. Furthermore, we identified SOX2/FOXA1 co-bound genes such as *UGT2B11* (**Figure 6C**), a gene implicated in production of intratumoral androgens and regulated by the NEPC-promoting transcription factor ONECUT2^66^.

We also identified SOX2/FOXA1 peaks near the *ROR1* gene (**Figure 6D**). The ROR family of receptor tyrosine kinases have been shown to mediate lineage plasticity in PCa^59^, and ROR1 may act through an autocrine role, given that its ligand is *WNT5A*^58,67^, another SOX2/FOXA1 co-bound gene in NEPC (**Figure 4L**). We show that *ROR1* transcript levels are highest in our NEPC cell lines compared to adenocarcinoma, correlating with the expression of *SOX2* itself (**Figure S5G**).

In previous publications, we found that SOX2^KO^ CWR-R1 cells display altered metabolism, decreased proliferation, and increased sensitivity to AR-targeted therapies^25,26^. The transcript levels of genes near SOX2/FOXA1 co-bound regions were downregulated after SOX2 knockout (SOX2^KO^) in CWR-R1 cells^25^, as well as *FOXA1*, which is itself a putative SOX2/FOXA1 gene target (**Figures 6E and S5H**). Following SOX2^KO^, AR is upregulated along with AR-responsive *COLCA1*, but not the neuroendocrine tumor-associated gene *COLCA2*, which is in line with previously published data^25,64^ (**Figure 6E**). In addition, we find that SOX2^KO^ results in a dramatic reduction of estrogen-related receptor gamma (*ESRRG*) and NOTCH-regulated ankyrin repeat protein (*NRARP*) transcripts, whose genes both have strong SOX2/FOXA1 co-binding in both CWR-R1 and NEPC (**Figures 6E and S5I-J**). Importantly, **Figure 6E** implies that SOX2 directly regulates the expression of key genes that both mediate ARSI resistance and facilitate the transition from adenocarcinoma to the lethal NEPC phenotype.

Looking at the putative SOX2-FOXA1 gene targets in CWR-R1, AR often binds with SOX2 and FOXA1 regardless of hormone manipulation (**Figure 6F-G**). The raw DNA binding peaks in **Figures 6F-G** suggest that SOX2 may mediate stabilization of this transcriptional complex; this may occur through a pioneering role with FOXA1 to promote activation of these genes with either full-length AR or ligand-independent AR-V7. When looking at which genes map near these conserved binding peaks (TSS within 10kb of the ChIP-seq peak) (**Figure 6H**), genes implicated in membrane lipid metabolism and estrogen metabolism are enriched (**Figure 6I**), suggesting that these genes are important in driving the ARSI-resistance and proliferation we have previously shown SOX2 mediates^15,25,26^.

We then chose to focus on the putative SOX2/FOXA1 target gene *ROR1*, given that its signaling is ligand-activated by WNT5a to promote the non-canonical oncogenic Wnt pathway in metastatic CRPC^67^, and that there are ongoing clinical efforts to ablate ROR1 activity in CRPC^68^. In addition, ROR2, the WNT5a-activated heterooligomeric partner of ROR1, is an important upstream regulator of *ASCL1* that promotes enzalutamide resistance, lineage plasticity, and the NEPC phenotype^59^. As was seen following SOX2^KO^ in CWR-R1 cells (**Figure 6E**), SOX2 siRNA knockdown in H660 leads to a significant decrease in *ROR1* expression (**Figure 6J**). Given the results from **Figure 6E and 6J**, which establish *ROR1* is a *bona fide* SOX2 target gene in both adenocarcinoma and NEPC, we then sought to use its gene sequence to computationally model the SOX2-FOXA1 protein interaction on DNA containing their respective FOX and HMG-box motifs (**Figure 6K and 6L**). *In silico* structure models of the SOX2/FOXA1 heterodimer at the *ROR1* peak summit demonstrate a substantial contact interface between the two proteins (**Figure S6A-B**). The SOX2 HMG-box domain engages with the DNA minor grove, alongside the FOXA1 forkhead domain (FKHD) binding the same face of the DNA downstream (**Figure 6M**). We also predict three-dimensional looping of the DNA by the SOX2-FOXA1 heterodimer (**Figure 6N**), consistent with the mechanism in which SOX2 cooperates with OCT3/4 near nucleosomes in hESCs^35^. Taken together, these data underscore the functional importance of a strong SOX2-FOXA1 interaction, providing key insights into how their coordinated behavior mediates target gene expression, ARSI-resistance, and lineage plasticity in PCa.

### *SOX2* expression is highest in AR-low patient tumors, especially in FGFR-driven Double Negative Prostate Cancer (DNPC) and displays a unique cistrome in cancers compared to normal tissues

Since SOX2 is AR-repressed, yet still expressed in CRPC, we probed published single-cell RNA-sequencing (scRNA-seq) datasets of heavily treated prostate cancer patient tumors (**Figure 7A)**^69–71^. We stratified these patient samples by subtype (**Figure 7B**) and by NEPC signature score (**Figure 7C**). In addition, we scored the samples based on *AR*, *FOXA1*, and *SOX2* expression (**Figure 7D-F**). We find that tumors with the highest *AR* expression unsurprisingly have the least *SOX2* expression (**Figure 7F**), with FOXA1 being mostly invariant (**Figure 7E and 7G**). Interestingly, one tumor (HMP08) is representative of a “double-negative” prostate cancer (DNPC), a tumor that is neither NEPC marker nor AR-positive^72,73^. HMP08 has the highest *SOX2* expression (**Figure 7F**), while one neuroendocrine tumor (HMP17) has lower expression of AR, FOXA1, and SOX2 (**Figure 7G**) and is high for *NEUROD1* expression, but not *ASCL1* expression (**Figure S7A**) ^74^. These data fit our model, as *ASCL1* seems to be a SOX2-FOXA1 target (**Figure 4H**), its expression correlates with *SOX2* in patient samples (**Figure S7B-D**), and it has an established relationship with the ROR family proteins^59^. Furthermore, in an *in vivo* temporal model of NEPC trans-differentiation [Pan-small cell neuroendocrine cancer (PARCB)]^56^, *SOX2* expression is positively correlated with *ASCL1* expression (**Figure S7E**), is not correlated with *NEUROD1* expression (**Figure S7F**), and unsurprisingly, is negatively-correlated with *AR* expression (**Figure S7G**). Interestingly, previous studies have shown that DNPC tumors are associated with FGF signaling^72,73^, consistent with our published findings identifying *FGF5* as a SOX2 target^15^ (**Figure 6E, Figure S5D).**

**Figure 7.**
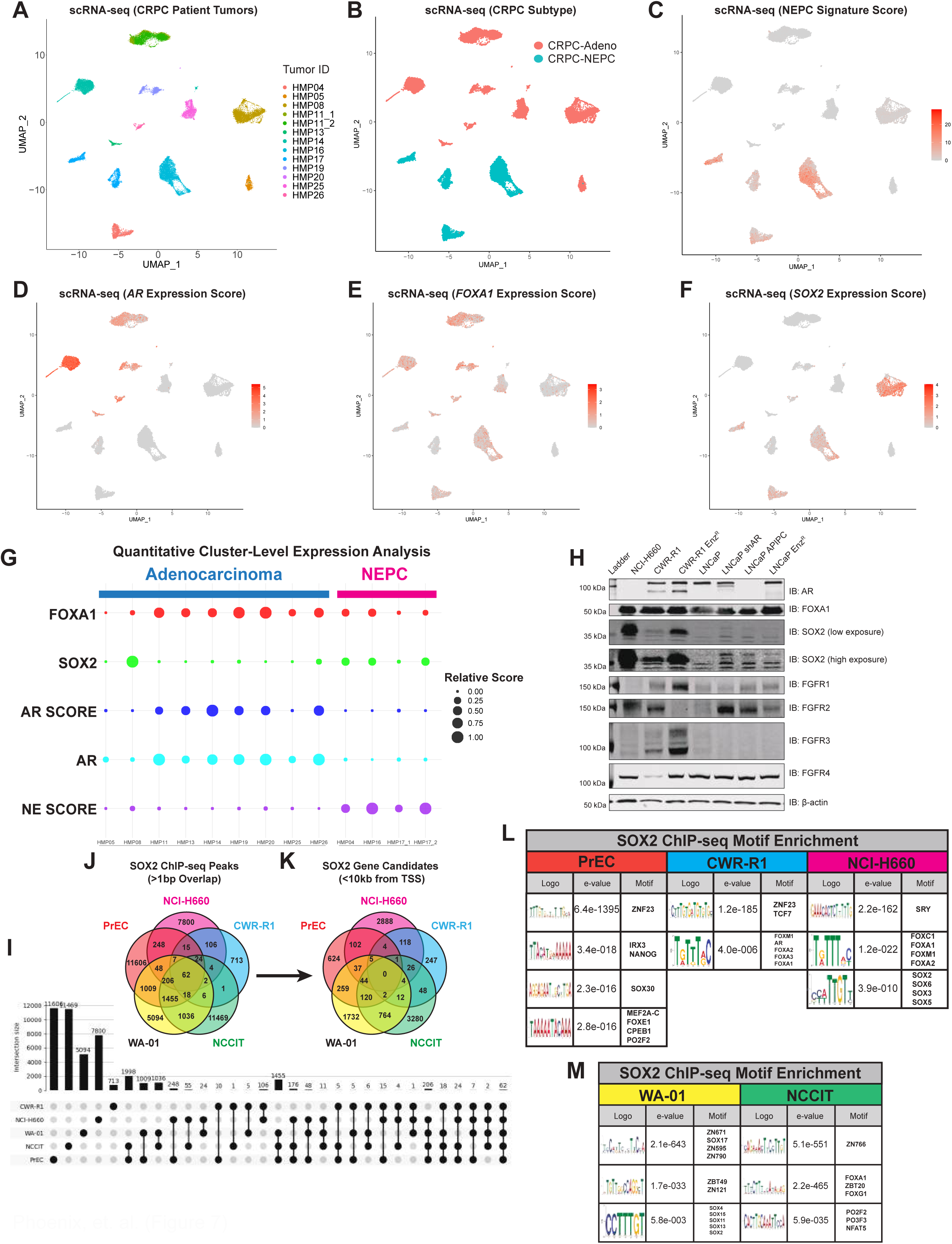
*SOX2* expression is highest in AR-low patient tumors, especially in FGFR-driven Double Negative Prostate Cancer (DNPC) and displays a unique cistrome in cancer compared to normal tissues. **A)** UMAP visualization of prostate cancer cell populations from 13 castration-resistant patients from the Human Metastatic Prostate (HMP) dataset. **B)** UMAP visualization of the same 13 patients categorized by histological type: adenocarcinoma and NEPC. **C)** NEPC signature analysis applied to CRPC patients. **D-F)** Expression analyses of *AR*, *FOXA1*, and *SOX2* across the same UMAP coordinates **G)** Expression levels of genes (*AR*, *FOXA1*, *SOX2*) and pathway signatures (AR, NEPC scores) are shown through a bubble plot as organized by sample. **H)** Western blot panels of malignant prostate cancer cells for AR, FOXA1, SOX2, and the FGFR-family of proteins. Blots for β-actin were used as an internal loading control. **I)** UpSet plot of SOX2 ChIP-seq peaks in CWR-R1, NCI-H660, PrEC, WA-01 and NCCIT cells. Shared regions represent called peaks with >= 1bp of overlap. **J)** Venn Diagram SOX2 ChIP-seq peaks in CWR-R1, NCI-H660, PrEC, WA-01 and NCCIT cells. Shared regions represent called peaks with >= 1bp of overlap. **K)** Venn diagram of SOX2 potential target genes. SOX2 ChIP-seq peaks from all cell lines were analyzed using Cistrome-GO to identify genes that SOX2 binding peaks less than 10kb from a TSS and an adjusted regulatory potential (RP) score of >0.01. We find zero genes that are commonly bound by SOX2 across the different cell contexts. **L)** Binding motif enrichment near SOX2 peaks in PrEC, CWR-R1, and NCI-H660 cells. Both MEME and STREME motif discovery analyses were included. Full lists of motifs discovered can be found in Supplementary Table 1. **M)** Binding motif enrichment near SOX2 peaks in pluripotent WA-01 and NCCIT cells. Both MEME and STREME motif discovery analyses were included. Full lists of motifs discovered can be found in **Supplementary Table 1**.

We then compared levels of SOX2 and FOXA1 protein among cell line models of DNPC, NEPC, and ARSI-resistance: LNCaP APIPC (the cell line model of DNPC), their parental shAR derivatives (LNCaP shAR), the isogenic SOX2-negative LNCaP cells, their enzalutamide-resistant counterpart LNCaP-Enz^R^, SOX2-expressing H660 (NEPC), and CWR-R1 and CWR-R1 Enz^R^ adenocarcinoma cell lines. *SOX2* expression is present in the LNCaP APIPC and LNCaP shAR cell lines, on par with the expression in the LNCaP-Enz^R^ (**Figure 7H**). However, by comparison, SOX2 levels are lower than in the CWR-R1 and H660 cells (**Figure 7H**). We also assessed levels of the FGFR family proteins in our cell lines and found expression levels to be mixed. Notably, we observe higher FGFR2 levels in CWR-R1 and the LNCaP cell lines that express SOX2 (**Figure 7H**). In addition, we note that FGFR1 and FGFR3 levels are highest in the CWR-R1-Enz^R^ and H660 (**Figure 7H**). Perhaps, this could suggest an upstream role of these proteins in promoting *SOX2* expression; however, outside of FGF signaling, SOX2 has been implicated in basal epithelial tumor expression in DNPC^75^.

We find a positive correlation of SOX2 with basal and neuroendocrine signature (**Figure S7H-I**) and no correlation with luminal signature or with AR activity in PCa (**Figures S7J-K).** When looking specifically at which transcripts are associated with a basal transcriptional signature in CRPC, we find genes like *SOX2* and the FGFR family to be positively correlated, and genes like *FOXA1*, *AR*, and *NKX3-1* to be negatively correlated (**Figure S7L**).

We previously identified an important role of SOX2 in AR-negative, normal basal prostate epithelial cells (PrEC)^15,76^, and as such, performed ChIP-seq on *SOX2*-expressing, AR-negative PrEC derived as a primary culture from normal prostate tissue grown in keratinocyte serum-free media^77^. We compared the DNA binding profile of SOX2 in PrECs to that in H660, CWR-R1, pluripotent hESCs (WA-01), and its transformed counterpart human embryonal carcinoma cell line NCCIT^78^ (**Figure 7I**). There is very little overlap between peaks (**Figure 7J**) and zero overlap between candidate target genes within 10kb of SOX2 peaks across all cell lines (**Figure 7K**). When looking at the DNA motifs enriched near SOX2 ChIP-seq peaks in these cell lines, we do not see enrichment of FOXA1 motifs in PrEC (**Figure 7L-M**), nor in the canonical SOX2 context of hESC (WA01). However, in the pluripotent-like embryonal carcinoma cells (NCCIT), which have high OCT3/4 expression, there is some enrichment of a FOXA1 motif (**Figure 7M**). These data together suggest that the SOX2-FOXA1 interaction may be unique to cancer cells in adults, providing a potential therapeutic target in blocking the interaction or downstream effectors in AR-positive adenocarcinoma or NEPC.

## Discussion

Although SOX2 has a well-defined role as a stem-cell transcription factor and is canonically thought of as a Yamanaka factor^16^ governing both pluripotent and adult stem cell identity^30^, it is also a potent oncogene outside of the stem-cell context. It has been implicated as an oncogene in at least 25 different tumor types, including esophageal, lung, brain, breast, cervical, ovarian, and importantly, prostate cancer^79^. However, in tumors, SOX2 does not seem to necessarily induce a stem-like phenotype and is not always expressed with stem-cell markers, such as CD133 or even other Yamanaka transcription factors^15,79^. SOX2 has been implicated as a driver of ARSI resistance and lineage plasticity in prostate cancer^15,26,80,81^, yet it cannot act alone as it is an obligate heterodimer^29^. Here, we show that *SOX2* expression is necessary when it is expressed in both AR-positive adenocarcinoma and AR-negative NEPC cells (**Figure 1**). However, despite the lack of canonical SOX2 partners, in this study we show that SOX2 instead utilizes FOXA1 as its heteromeric partner in a cancer-specific manner to drive survival and proliferation of both prostate adenocarcinoma and NEPC.

Similarly to what has been previously ascribed to Nanog^82,83^, SOX2 binds to FOXA1 and engages a subset of AR targets that may mediate ARSI and stabilizes AR DNA binding near SOX2 with FOXA1. This is of note because *Nanog* is not concurrently expressed with *SOX2* and is positively upregulated by AR, unlike SOX2^84^. *SOX2* expression is negatively correlated with AR, as we have previously shown that *SOX2* is transcriptionally repressed by AR.^15^ However, co-expression of both *SOX2* and *AR* still occurs in patient tumors, and their expression is uniform across cells within tumors and not limited to a small or stem-cell like population^15^. Our data also suggest that AR and SOX2 do not directly interact, and in fact compete for binding to FOXA1 (**Figure 2**). In CWR-R1, SOX2 facilitates binding near genes regulated by AR and FOXA1 to promote a subset of pro-proliferative genes (**Figure 3**) and those that are known to mediate ARSI, such as steroid and androgen metabolism genes (**Figure 6**).

Given that both SOX2^85^ and FOXA1^5^ can serve as pioneering factors, SOX2 may serve to open chromatin AR with other AR cofactors at these sites in the absence of androgens or during antagonism. In particular, the AR splice variant AR-V7, which requires no ligand to bind DNA and is highly expressed in CWR-R1 and may bind preferentially at these sites^86^. Furthermore, both AR^87^ and SOX2^85^ have been shown to form phase-separated condensates that can host a variety of proteins on the DNA, it stands to reason that they may exist in a broader complex together with FOXA1 within the same cells. Previously, we identified NKX3.1, FOXA1 and AR together in a transcriptional context via similar techniques^42^, and interestingly, NKX3.1 has been shown to be able to replace OCT4’s role as a Yamanaka factor during pluripotent stem cell reprogramming^88^. Perhaps NKX3.1 is important in mediating AR and SOX2-FOXA1 interactions in this complex. However, this all requires more investigation, given that assays performed in bulk cell populations may obscure differences in expression in single cells, in cellular subpopulations, or among cells in different phases of the cell cycle.

SOX2 overlaps with FOXA1 binding nearly 90% of the time in both adenocarcinoma cells and NEPC. In NEPC, SOX2 binds near many classic markers of NEPC, produces a neuronal precursor signature consistent with an NEPC gene expression profile, and is positively correlated with expression in patient samples (**Figures S5E and S5F)**. Of the many SOX2/FOXA1 co-bound genes, we find particularly interesting three members of the non-canonical Wnt-signaling pathway, *ROR1*, *ROR2*, and *WNT5A* (**Figures 4L, S5E, and 6D**). Since SOX2 knockdown in H660 cells results in decreased *ROR1* transcript and SOX2 knockout leads to decreased *ROR1* and *WNT5A* mRNA in CWR-R1 (**Figure 6E and 6J**), these data suggest that SOX2 and FOXA1 may co-activate expression of *ROR1* and *WNT5A*, which in turn promotes the establishment of a feed-forward loop of non-canonical oncogenic Wnt-signaling in both CRPC and NEPC. That, in combination with ROR2 being upstream of ASCL1 in NEPC^59^, asserts this set of co-regulated SOX2/FOXA1 target genes as promising novel NEPC targets. Another SOX2/FOXA1 co-bound gene, *SEMA3C* (**Figure S5F**), is known to be upregulated specifically by pathogenic FOXA1 mutants^61^ with which we show SOX2 retains its ability to heterodimerize (**Figure S2D**). Furthermore, we previously identified that *SOX2* overexpression promotes castration-resistant growth of LAPC-4 cells^15^, which harbor an oncogenic C-terminal truncation of FOXA1 that promotes chromatin binding^31^. Together, this suggests that many oncogenic FOXA1 mutations can still bind with SOX2 and function transcriptionally.

In NEPC, other groups report that the POU/OCT family member BRN2 (*POU3F2*) binds SOX2 in NEPC models^28^. As we stated previously, NEPC is associated with gene expression normally found in the developing brain^47–49^. *SOX2* expression is essential for neural progenitor identity^89^ and neuronal development at different stages, not only by heterodimerizing with BRN2 in the developing brain^90^, but also PAX6 and ATRX^91^. BRN2 may bind with FOXA1 and SOX2 in a complex, or to SOX2 alone, as it may be responsible for the residual ∼10% of SOX2 binding peaks without FOXA1 (**Figure 4**). Furthermore, BRN2 has a known function in binding ASCL1 in neuronal precursors^92^. The genes SOX2 and FOXA1 co-regulate may be a co-opted from a developmental pathway normally found in neurons. Future work is necessary to assess how these transcriptional complexes come together and what genes they regulate in these different contexts.

Another interesting finding is that FOXA1 binding sites do not seem to be overtly enriched near SOX2 peaks in PrEC (**Figure 7L**), and *FOXA1* expression does not correlate with a basal transcriptional signature (**Figure S7L**). This suggests that SOX2 may interact with another protein besides FOXA1 in basal epithelial cells and basal-like DNPC tumors, given its correlation with a basal-cell signature in tumors. For instance, in lung squamous cell carcinoma (LUSC), SOX2 has been shown to heterodimerize with p63 instead of OCT3/4^93^. P63 is a basal cell marker we previously found co-expressed with SOX2 in basal-derived PrEC^15^. Furthermore, in the neural-like LUSC, SOX2 has been shown to bind BRN2 instead of p63,^94^ as it does in NEPC^28^. However, we did not identify a BRN2 or a p63 motif in our analysis of the SOX2 ChIP-seq in PrEC (**Supplemental Table 1**). Future work will determine the partner of SOX2 in both normal basal PrEC and DNPC contexts, and whether these two cell types utilize the same SOX2 partner, given transcriptional similarities^75^.

Finally, we suggest that the SOX2-FOXA1 interaction is both oncogenic and unique to cancer in adults. There is little evidence to support a SOX2-FOXA1 interaction in pluripotent hESCs or in normal basal PrEC context, however, we identified FOXA1 motifs enriched in the SOX2 ChIP-seq in NCCIT embryonal carcinoma cells (**Figure 7**). We propose that this presents a therapeutic opportunity in targeting the interaction; either directly through small molecules, peptide memetics, or through targeting downstream SOX2/FOXA1 effectors. In addition, we have shown that cell cycle genes are likely regulated by the complex and have already shown utility in targeting SOX2-positive prostate cells with anti-proliferative drugs such as WEE1 inhibitors^26^. Altogether, here we establish SOX2 and FOXA1 as heteromeric transcriptional binding partners that work together to co-regulate a subset of genes which promote PCa cell proliferation and confer lineage plasticity, regardless of AR status, in both CRPC and NEPC. Our future efforts will be dedicated to disrupting the SOX2-FOXA1 interaction directly or drugging the downstream signaling pathways. In late-stage prostate cancer, which currently has very few effective therapeutic options, innovative approaches beyond AR-targeted therapies are desperately needed; here, we assert the SOX2-FOXA1 interaction and its downstream signaling pathways as mechanistic drivers of resistance to AR-targeted therapies and aggressive, lineage plastic PCa phenotypes.

## Supporting information

Key Resources Table

## SUPPLEMENTAL FIGURES

**Figure S1.**
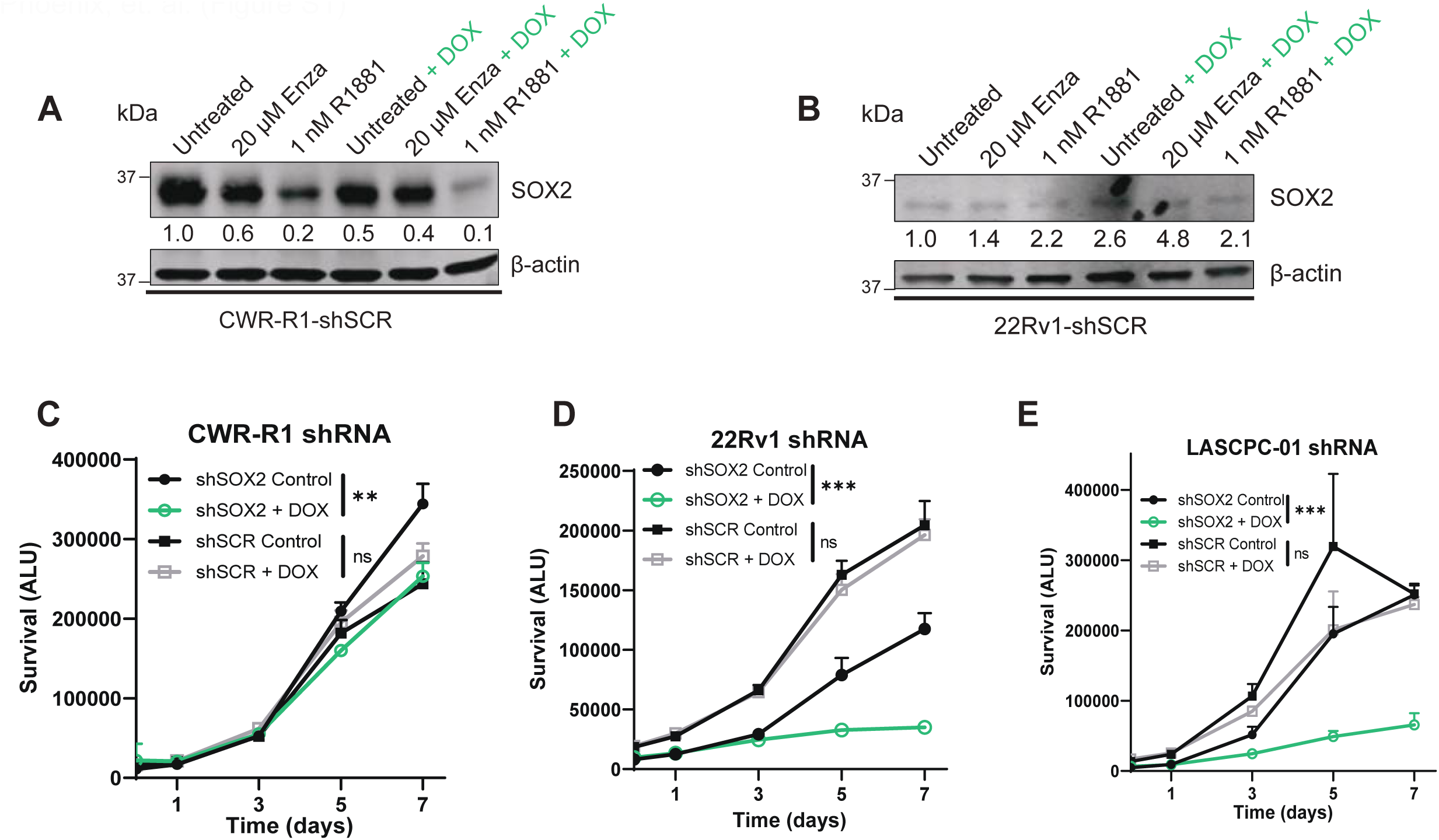
SOX2 knockdown invokes a proliferation defect in both CRPC adenocarcinoma and NEPC cells. **A-B)** Western blots show SOX2 protein levels are unaffected by induction of scrambled shRNA at 48h in CWR-R1 and 22Rv1 adenocarcinoma cells. Protein bands were quantified using Empiria Studio. **C-E)** Growth curves of CWR-R1, 22Rv1, and LASCPC-01 PCa cells across 7 days after dox-induction of shRNAs targeting either *SOX2* or scrambled control. Data points represent mean luminescence +/- SD.

**Figure S2.**
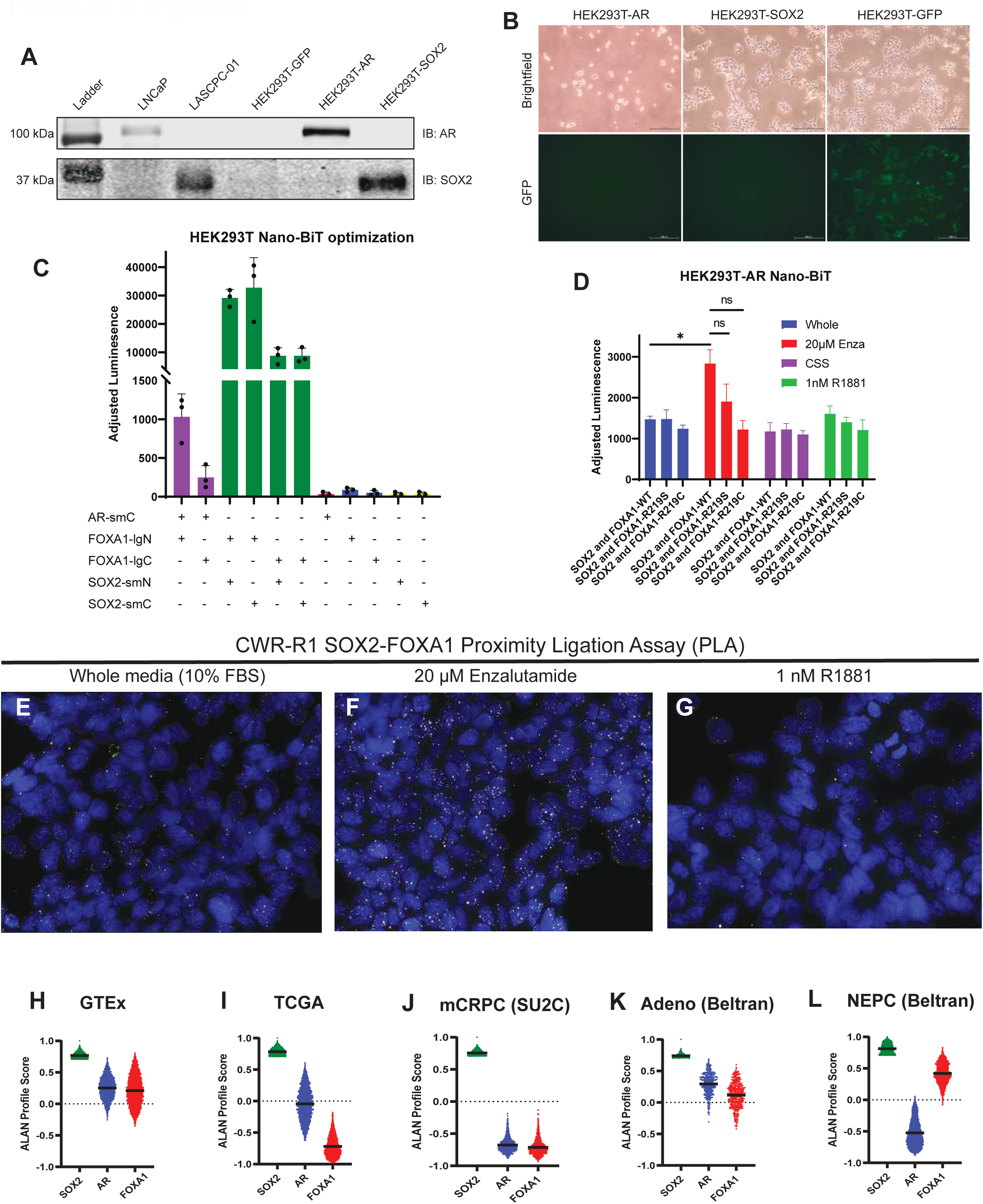
Lentiviral AR, SOX2, and GFP overexpression and optimization of Nano-BiT assays in HEK293T cells. **A)** Western blots show successful overexpression of AR and SOX2 protein after lentiviral transduction of HEK293T cells. LNCaP and LASCPC-01 cells are included as controls. **B)** Brightfield (top row) and GFP (bottom row) microscopy images of the lentivirus-transduced, puromycin-selected HEK293T cells. **C)** Split nano-luciferase complementation reporter assay (Nano-BiT) optimization shows a specific interaction between SOX2 and FOXA1 in HEK293T cells, regardless of the terminal tag location. Tagged AR and FOXA1 were shown to interact as a positive control, and single-BiT controls were included as a negative control. Data points represent mean luminescence +/- SD. **D)** Nano-BiT in AR-overexpressing HEK293T cells shows no change in SOX2 interaction with clinically relevant FOXA1 mutants. Data points represent mean luminescence +/- SD.

**Figure S3.**
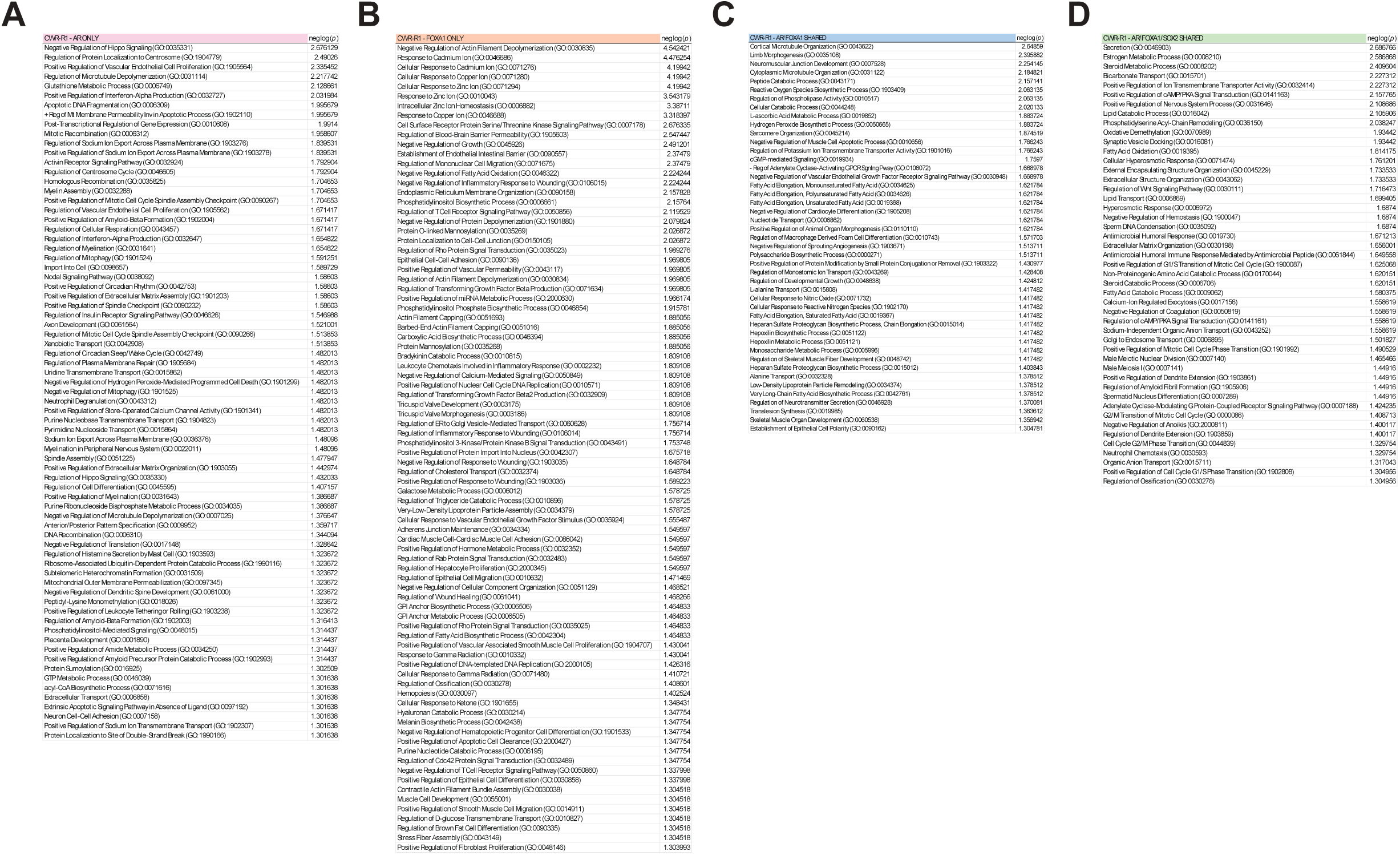
Full list of AR, FOXA1, and SOX2 binding sites’ Gene Ontology (GO) Biological Processes enrichment in CWR-R1 adenocarcinoma cells. **A)** Full list of significantly enriched (*p* <= 0.05) GO Biological Processes for candidate genes of near AR binding sites alone in CWR- R1 cells. GO enrichment was performed using Enrichr; data shown as -log(*p*). **B)** Full list of significantly enriched (*p* <= 0.05) GO Biological Processes for candidate genes of near FOXA1 binding sites alone in CWR-R1 cells. GO enrichment was performed using Enrichr; data shown as -log(*p*). **C)** Full list of significantly enriched (*p* <= 0.05) GO Biological Processes for candidate genes of near AR/FOXA1 co-bound sites without SOX2 in CWR-R1 cells. GO enrichment was performed using Enrichr; data shown as -log(*p*). **D)** Full list of significantly enriched (*p* <= 0.05) GO Biological Processes for candidate genes of near AR/SOX2/FOXA1 co-bound sites in CWR- R1 cells. GO enrichment was performed using Enrichr; data shown as -log(*p*).

**Figure S4.**
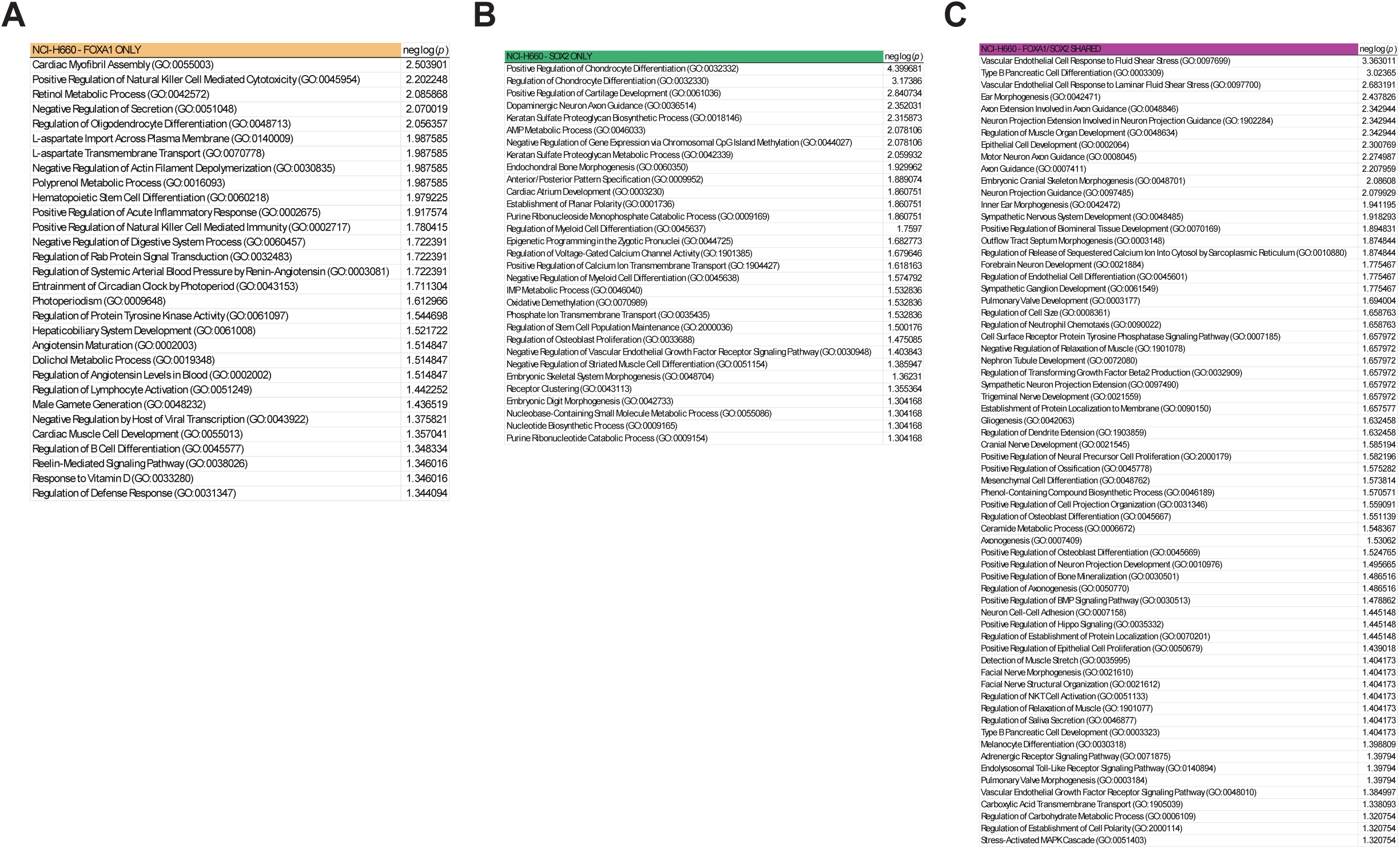
Full list of SOX2 and FOXA1 binding sites’ Gene Ontology (GO) Biological Processes enrichment in NCI-H660 NEPC cells. **A)** Full list of significantly enriched (*p* <= 0.05) GO Biological Processes for candidate genes of near FOXA1 binding sites alone in NCI-H660 NEPC cells. GO enrichment was performed using Enrichr; data shown as -log(*p*). **B)** Full list of significantly enriched (*p* <= 0.05) GO Biological Processes for candidate genes of near SOX2 binding sites alone in NCI-H660 NEPC cells. GO enrichment was performed using Enrichr; data shown as -log(*p*). **C)** Full list of significantly enriched (*p* <= 0.05) GO Biological Processes for candidate genes of near SOX2/FOXA1 co-bound sites a in NCI-H660 NEPC cells. GO enrichment was performed using Enrichr; data shown as -log(*p*).

**Figure S5.**
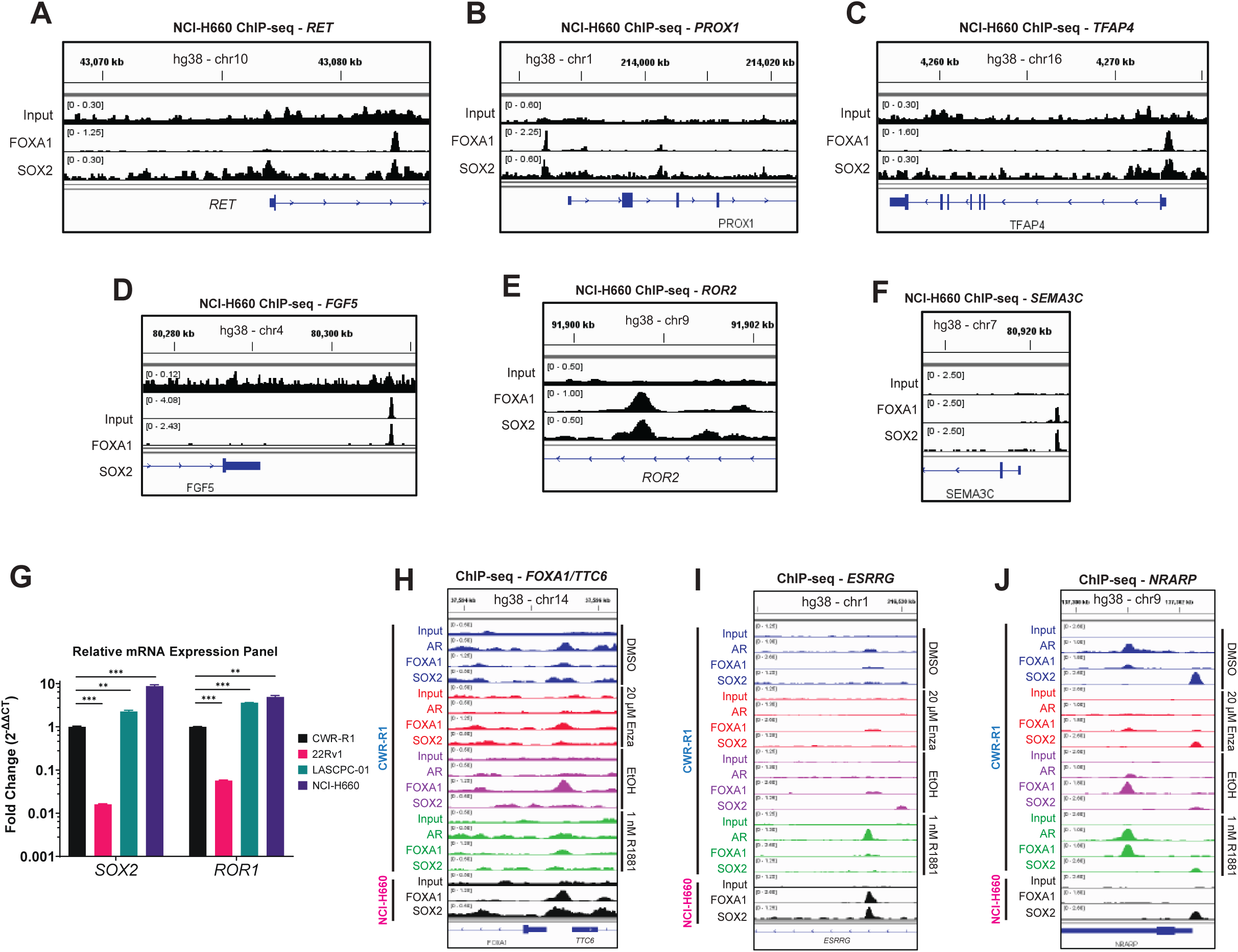
Candidate SOX2 target genes in CRPC and NEPC cell line models. **A-F)** Track plots of SOX2 and FOXA1 ChIP-seq peaks near oncogenic/NEPC genes in NCI-H660. Peaks visualized using IGV and mapped to the human hg38 chromosome build. **G)** Relative mRNA expression of *SOX2* and *ROR1* across PCa cell lines normalized to β-actin. Data are represented as the fold change (2^-ΔΔCT^) ± SEM. **H-J)** Track plots of AR, FOXA1, and SOX2 co-bound genes in CWR-R1 and NCI-H660. Peaks visualized using IGV and mapped to the human hg38 chromosome build.

**Figure S6.**
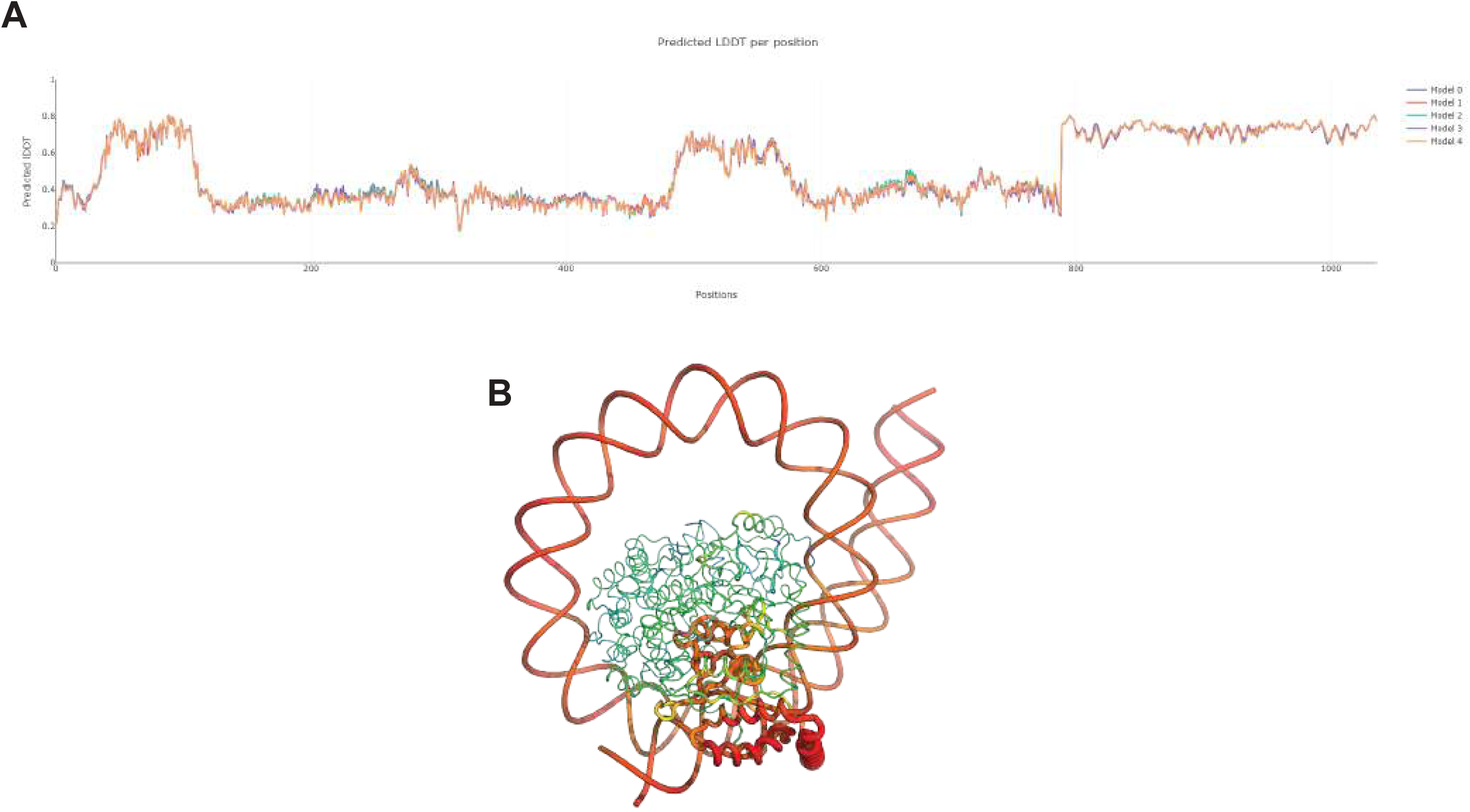
Confidence for the predicted FOXA1 and SOX2 complex on DNA within the *ROR1* gene. **A)** Predicted local difference distance test (LDDT) per position. Scores between 0.7 and 0.9 indicate confident regions of prediction. **B)** Heatmap of predicted LDDT per position whereby regions of higher confidence are shown as hotter colors with wide tubes and lower confidence scores are shown in thin cooler colors.

**Figure S7.**
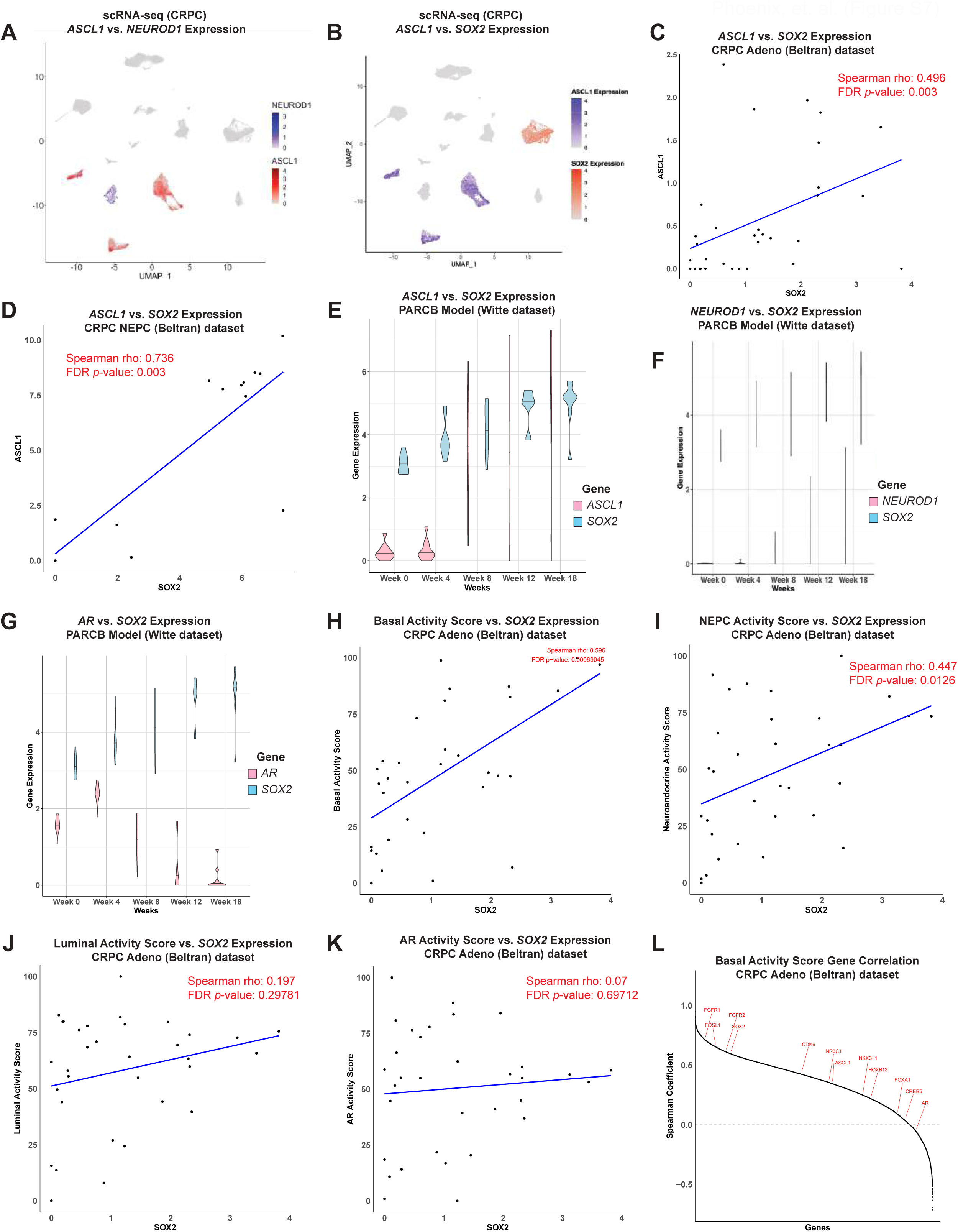
*SOX2* expression is correlated with oncogenic transcriptional signatures and other key mediators of lineage plasticity and the NEPC phenotype. **A)** Expression analyses of *ASCL1* and *NEUROD1* across the same CRPC patient UMAP coordinates from the HMP dataset in **Figure 7**. **B)** Expression analyses of *SOX2* and *ASCL1* across the same CRPC patient UMAP coordinates from the HMP dataset in **Figure 7. C)** Scatter plot showing the relationship between *SOX2* and *ASCL1* expression in the Beltran (Adeno)^40^ dataset. **D)** Scatter plot showing the relationship between *SOX2* and *ASCL1* expression in the Beltran (NEPC)^41^ dataset. **E)** Violin plot showing gene expression changes over time for *ASCL1* alongside *SOX2*. Violin plot expression levels were obtained from the *in vivo* temporal model of NEPC trans-differentiation [Pan-small cell neuroendocrine cancer (PARCB)]^56^ and are displayed across different time points, highlighting temporal patterns and co-expression relationships between target genes and *SOX2*. **F)** Violin plot from the PARCB model showing gene expression changes over time for *NEUROD1* alongside *SOX2*. **G)** Violin plot from the PARCB model showing gene expression changes over time for *AR* alongside *SOX2*. **H)** Scatter plot showing the relationship between *SOX2* expression and Basal Activity Score the Beltran (Adeno) dataset. **I)** Scatter plot showing the relationship between *SOX2* expression and Neuroendocrine Activity Score the Beltran (Adeno) dataset. **J)** Scatter plot showing the relationship between *SOX2* expression and Luminal Activity Score the Beltran (Adeno) dataset. **K)** Scatter plot showing the relationship between *SOX2* expression and AR Activity Score the Beltran (Adeno) dataset. **L)** Graph ranking Spearman correlation coefficients for select genes with respect to Basal Activity Score profiles in the Beltran (Adeno) dataset. Genes like *SOX2* and *FGFR1* show significant positive correlations, whereas genes such as AR, FOXA1, and NKX3-1 are negatively correlated with Basal Activity Score.

## Materials and Methods

### Cell Culture

Unless otherwise stated, all cell lines were purchased from American Type Culture Collection (ATCC) and incubated at 37 ^ͦ^ C with 5% CO_2_. CWR-R1, LNCaP, NCCIT, and 22Rv1 were grown using 10% FBS (BioWest Cat: S1480) RPMI-1640 with ATCC modification (ATCC, Cat: 30-2001). VCaP and HEK293T cells were cultured in 10% FBS DMEM (ATCC, Cat: 30-2002). NCI-H660 and LASCPC-01 were grown using 5% HITES (5% FBS, 10nM β-estradiol (Sigma-Aldrich, Cat: E2578), 10nM hydrocortisone (Sigma-Aldrich, Cat: H0888), 1% ITS-G (Gibco, Cat: 41400-045)) RPMI-1640 with ATCC modification. PNT-2, PC-3, AND DU-145 cells were cultured in 10% FBS RPMI-1640 (Gibco, Cat: 11875093). LNCaP-APIPC and LNCaP-shAR controls were generously supplied by Dr. Peter Nelson at UW Fred Hutchinson Cancer Center. LNCaP-APIPC were maintained in Phenol-red-free RPMI-1640 (5% CSS, 2.5 µg/mL blasticidin, 1 µg/ml puromycin, 25 µg/ml Zeocin, 1 µg/mL doxycycline). LNCaP-shAR were maintained in Phenol-red free RPMI-1640 (5% FBS, 2.5 µg/mL blasticidin, 1 µg/ml puromycin, 25 µg/ml Zeocin). All other cell lines, including enzalutamide resistant counterparts, were cultured as previously described^27^. Non-malignant PrEC cultures were established from fresh human prostate tissues acquired from surgical specimens as previously described^15^. These tissues were acquired under an expedited protocol approved by the University of Chicago Institutional Review Board (IRB). Tissue samples were managed by the University of Chicago Human Tissue Resource Center core facility; the need for patient consent was waived as acquired samples were de-identified. Dissociation of prostate tissue and growth of epithelial cells has been previously described^15^ and the same methods were used to establish matched PrEC cultures. Cultures were grown using Keratinocyte Serum-Free Defined media supplemented with growth factors (ScienCell, Cat: 2101). For our experiments, all cultures were analyzed at or before their fourth passage. Unless otherwise stated, cells treated with enzalutamide (Selleckchem, Cat: S1250) were treated at a final concentration of 20 μM and cells treated with R1881 (Sigma-Aldrich, Cat: R0908) were treated at a final concentration of 1 nM.

### Western Blotting

Whole cell lysates were collected from cells in cold RIPA buffer (150 mM sodium chloride (Sigma-Aldrich, Cat: S9888), 1.0% Igepal CA-630 (Sigma-Aldrich, Cat: I8896), 0.5% sodium deoxycholate (Sigma-Aldrich, Cat: D6750), 0.1% SDS (Sigma-Aldrich, Cat: 436143), 50 mM Tris (Sigma-Aldrich, Cat: 648311, pH 8.0) containing 1X Protease/Phosphatase inhibitor cocktail (MP Biomedicals, Cat: 08W00017). Lysates were then homogenized using a probe sonicator (ThermoFisher, Cat: FB-120). Protein levels were quantified using a Pierce BCA kit (ThermoFisher, Cat: 23225) and 20μg of protein was loaded per lane into 10% SDS-PAGE gels. After transferring to nitrocellulose membranes (Bio-Rad, Cat: 1620115), blotting was performed with primary antibodies [AR - rabbit monoclonal anti-AR(XP), Cell Signaling (Cat: 3579) | FOXA1 - rabbit polyclonal anti-FOXA1, ThermoFisher (Cat: PA5-27157) or mouse monoclonal anti-FOXA1, ThermoFisher (Cat: MA1-091) | SOX2 - rabbit monoclonal anti-SOX2(XP) (Cell Signaling, Cat: 3579) or rabbit polyclonal anti-SOX2 (Millipore, Cat: AB5603) | FGFR1(XP) rabbit monoclonal antibody (Cell Signaling, Cat: 9740) | FGFR2 rabbit monoclonal antibody (Cell Signaling, Cat: 23328) | FGFR3 rabbit monoclonal antibody (Cell Signaling, Cat: 4574) | FGFR4 rat monoclonal antibody (Novus Biologicals, Cat: 240929) | Mouse monoclonal anti-β-actin (Millipore, Cat: A1978)]. Subsequent secondary antibody incubations (LICOR, Cat: 926-68072 | LICOR, Cat: 926-32213) were performed and blots were visualized using the LICOR Odyssey DLx system (LICOR, Cat: 9142-00).

### Generation of dox-inducible shRNA cell lines

STBL3 *E. coli* bacteria in glycerol stabs containing either Tet-pLKO-puro-shSOX2 (Addgene plasmid #47540, RRID:Addgene_47540) or Tet-pLKO-puro-shScrambled (Addgene plasmid #47541, RRID:Addgene_47541) were streaked for isolation 1.2% LB agar plates containing 100 μg/mL ampicillin (GoldBio, Cat: A-301-10) and incubated at 37 ^ͦ^ C for 16 hours (both plasmids were a gift from Charles Rudin^95^). Isolated colonies were picked and grown at 37^ͦ^ ^C^ for 16 hours in 5mL LB/ampicillin broth. We then isolated and purified our expression vector plasmid(s) from the bacteria using the GeneJET Miniprep kit (ThermoFisher, Cat: K0502) and verified concentrations/purity via NanoDrop (ThermoFisher, Cat: ND-2000C). All plasmids were stored at −20 ^ͦ^ C unless otherwise stated. We co-transfected viral packaging plasmid pCMV-dR8.2 (Addgene plasmid #8455) and envelope plasmid pVSV-G (Addgene plasmid #8454) (both lentivirus plasmids were a gift from Bob Weinberg)^96^ along with our validated expression vector (Tet-pLKO-puro-shScrambled or Tet-pLKO-puro-shSOX2) into HEK293T feeder cells grown in 10% FBS DMEM in 6-well plates using the ThermoFisher Lipofectamine 3000 kit (Invitrogen, Cat: L3000-015). For the co-transfection, we used 3 μg each expression vector, 1.5 μg packaging plasmid pCMV-dR8.2 per well, 0.75 μg envelope plasmid pVSV-G per well, and 2 μL P3000 reagent (from the Lipofectamine 300 kit) per μg of total plasmids transfected. Following co-transfection, the HEK293T cells were incubated overnight 37 ^ͦ^ C (5% CO_2_) for 24 hours. The following day, we harvested lentivirus directly from the HEK293T 6-well plates and performed 2 rounds of 0.45 μm syringe filtration (ThermoFisher, Cat: 725-2545) and centrifugation to remove HEK293T cells from suspension. After filtration, we combined the lentivirus (2mL) with 2 x 10^6^ of our PCa cells of interest (CWR-R1, 22Rv1, and LASCPC-01) in 2mL whole media suspension (1:1 virus/cell ratio by volume) plus final 8 μg/mL polybrene and plated them in T25 flasks which were incubated overnight 37 ^ͦ^ C (5% CO_2_) for 24 hours. The following day, we performed an additional round of lentivirus infection on the PCa cells by replacing the media with 1:1 virus/whole media ratio (by volume) plus final 8 μg/mL polybrene (Sigma-Aldrich, Cat: TR-1003-G) in the same T25 flask followed by incubation overnight at 37 ^ͦ^ C (5% CO_2_) for an additional 24 hours. After 48 hours, we replaced the media of our PCa cells with whole media supplemented with puromycin (Gibco, Cat: A11138-03) at a final concentration of 1 μg/mL to select for cells which successfully stably integrated our tet-inducible shRNA expression constructs. Cells were continued in puromycin for 2 passages until the mock transfected cells died from negative selection in puromycin.

### siRNA transfection

Parental NCI-H660 cells were plated in both 6-well (5 x 10^5^ cells/well) and 96-well plates (2000 cells/well) in 100 μL whole media and incubated at 37 ^ͦ^ C (5% CO_2_) for 24 hours. We then performed transfection of siRNAs targeting a non-specific sequence, 3 independent regions of the *SOX2* transcript, or *GAPDH* transcript [Silencer Select SOX2 siRNA #1 (Invitrogen, Cat: s13294) | Silencer Select SOX2 siRNA #2 (Invitrogen, Cat: s13295) | Silencer Select SOX2 siRNA #3 (Invitrogen, Cat: s13296) | Silencer Select GAPDH siRNA (Invitrogen, Cat: 4390849) | Silencer Select Negative Control #1 siRNA (Invitrogen, Cat: 4390843)] using the Lipofectamine RNAiMax kit (Invitrogen, Cat: 13778150) according to manufacturer’s protocol. 6 hours after the initial transfection, the 6- and 96-well plates were supplemented with 5% FBS HITES RPMI to a final volume of 2mL or 200μL, respectively, and incubated at 37 ^ͦ^ C (5% CO_2_) for a time course of 12 days. Protein was harvested after 48h in RIPA buffer and western blotting performed as stated previously.

### Cell Viability Assays

CWR-R1 (5000 cells/well), 22Rv1 (5000 cells/well), LASCPC-01 (2000 cells/well) dox-inducible shRNA cells were plated in 96-well plates and incubated at 37 ͦ C (5% CO2) for a time course of 7 days. 24 hours after plating, cells were treated with media containing 2μg/mL doxycyclineCell Titer-Glo 5min incubation then read on plate reader. Normalized to their own untreated (-DOX) controls at each time point. For cell viability experiments using the newly generated tet-inducible shRNA PCa cells, cells were plated in 96-well plates (Corning, Cat: 07-200-336) as follows: CWR-R1-shSCR and CWR-R1-shSOX2 (5000 cells/well), 22Rv1-shSCR and 22Rv1-shSOX2 (5000 cells/well), LASCPC-01-shSCR and LASCPC-01-shSOX2 (2000 cells/well), and incubated at 37 ^ͦ^ C (5% CO_2_) for 24 hours (n = 5 wells). Induction of shRNA was activated by replacing the cells’ media with their respective whole media containing final 2μg/mL doxycycline (Sigma-Aldrich, Cat: D3447). At the same time, the adenocarcinoma cells (CWR-R1 and 22Rv1) were treated with whole media containing either final 20μM enzalutamide or final 1nM R1881 to modulate AR activity. DOX media was replaced daily for the entire course, unless otherwise stated. For each cell line, protein lysate was harvested from 6-well plates 48h after treatment and subjected to western blotting as described previously. Cell viability assays were performed at day(s) 0, 1, 3, 5, and 7 for all cell lines using the Promega Cell Titer-Glo Kit (Promega, Cat: G7570) which was read using a BioTek Synergy H1 plate reader (Agilent, Cat: 11-120-533). Raw ALU (absolute luminescence units) for each condition (n = 5 wells) were normalized to each treatment conditions’ respective untreated control wells [ALU (+DOX) / ALU (-DOX)] at each time point and displayed as mean normalized survival ± SD. For cell viability experiments in NCI-H660 undergoing siRNA transfection, assays were performed at day(s) 0, 1, 3, 5, and 7 for all cell lines using the same protocol as other cell lines. In NCI-H660, raw luminescence values (ALU) for each condition (n = 6 wells) were recorded and displayed longitudinally as mean ALU (absolute luminescence units) +/- SD.

### Proximity Ligation Assays

CWR-R1 cells were seeded at 5 × 10^5^ cells per well in an 8-well glass chamber slide and incubated at 37°^C^ and 5% CO_2_ for 24 hours. Next, the cells were fixed with 4% paraformaldehyde in PBS (HyClone, Cat: SH3025602) for 20 min (on ice) with gentle shaking. Cells were then quenched with 50 mM ammonium chloride (Sigma-Aldrich, Cat: 213330) in PBS for 10 minutes. Cells were then washed 3 times for 5 min each at room temperature with PBS. Cells were then quickly rinsed with H_2_O to remove any salt. Plastic chambers were then removed from the slides and reaction areas were delimited with a grease pen on the boarder of each well. Next, we permeabilized the cells with 0.3% TritonX-100 (Sigma-Aldrich, Cat: X100) in PBS for 20 minutes at room temperature before being washed 3 times (5 min each) with PBS. We then performed the Duolink In Situ Red Starter Kit Mouse/Rabbit (Sigma-Aldrich, Cat: DUO92101) according to manufacturer’s protocol, starting at adding 1 drop of Duolink Blocking Solution to each well of the 8-well slide and incubating for 30 minutes at 37°^C^. Primary antibodies against proteins-of-interest used were rabbit monoclonal anti-SOX2(XP) (Cell Signaling, Cat: 3579) and rabbit polyclonal anti-FOXA1 (ThermoFisher, Cat: PA5-27157). Secondary-antibody-only controls as well as controls individually replacing each of the primary antibodies with normal Rabbit IgG (Cell Signaling, Cat: 3900) were used to ensure signals were not background. We imaged the slides on a Keyence BZX-800 fluorescence microscope (KEYENCE, Cat: BZ-X800) with a 60x-oil objective. All images were taken as 10-micron thick Z-stacks with a 0.4 μM step size and max projections of each stack were used for image analysis. Image analysis was done using Fiji^97^ to count the number of foci observed per individual nucleus. Foci that did not overlap with the nuclear DAPI signal were ignored.

### Co-Immunoprecipitations

Co-immunoprecipitation was performed on cells seeded at a density of 1.0 x 10^6^ cells per IP using a Dynabeads Protein G Immunoprecipitation Kit (ThermoFisher, Cat: 10007D). Cells were lysed with CellLytic Lysis Reagent (Sigma-Aldrich, Cat: C2978) and homogenized using a BioRuptor Pico Sonicator (Diagenode, Cat: B01080010). SOX2 was then immunoprecipitated with Dynabeads Protein G using beads conjugated to polyclonal goat anti-SOX2 antibody (R&D Systems, Cat: AF2018) or the species-matched goat IgG control antibody (R&D Systems, Cat: AB-108-C). FOXA1 was also immunoprecipitated with Dynabeads Protein G using beads conjugated to polyclonal rabbit anti-FOXA1 antibody (ThermoFisher, Cat: PA5-27157) or the species-matched rabbit IgG control antibody (Cell Signaling, Cat: 2729) Proteins were eluted in 2.5x Laemmli Buffer and loaded onto 10% SDS-PAGE gels and analyzed via western blot.

### Generation of AR, SOX2, and GFP-overexpressing HEK293T cells

We first transformed lentiviral overexpression plasmids LV105-AR, LV105-SOX2, and LV105-GFP (GeneCopoeia, Cat: EX-E2325-Lv105 | GeneCopoeia, Cat: EX-T2547-Lv105 | GeneCopoeia, Cat: eGFP-LV105) into DH5α *E. coli*, which were then streaked for isolation 1.2% LB agar plates containing 100 μg/mL ampicillin and incubated at 37 ^ͦ^ C for 16 hours. Colonies were picked and grown at 37 ^ͦ^ C for 16 hours in 5mL LB/ampicillin broth for 16 hours. We then isolated and purified our overexpression vector plasmid(s) from the bacteria using the GeneJET Miniprep kit and verified concentrations/purity via NanoDrop assay. All plasmids were stored at - 20 ^ͦ^ C unless otherwise stated. We co-transfected viral packaging (pCMV-dR8.2) and envelope (pVSV-G) plasmids along with our validated expression vector (LV-AR, LV-SOX2, LV-GFP) into HEK293T feeder cells grown in 10% FBS DMEM in 6-well plates using the ThermoFisher Lipofectamine 3000 kit. For the co-transfection, we used 3 μg each overexpression vector, 1.5 μg packaging plasmid pCMV-dR8.2 per well, 0.75 μg envelope plasmid pVSV-G per well, and 2 μL P3000 reagent per μg of total plasmids transfected. Following co-transfection, the HEK293T feeder cells were incubated overnight at 37 ^ͦ^ C (5% CO_2_) for 24 hours. The following day, we harvested lentivirus directly from the HEK293T 6-well plates and performed 2 rounds of 0.45 μm syringe filtration. After purification, we combined the lentivirus (2mL) with 2 x 10^6^ of parental HEK293T in 2mL whole media suspension (1:1 virus/cell ratio by volume) plus final 8 μg/mL polybrene and plated them in T25 flasks which were incubated overnight 37 ^ͦ^ C (5% CO_2_) for 24 hours. The following day, we performed an additional round of lentivirus infection on the parental HEK293T cells by replacing the media with 1:1 virus/whole media ratio (by volume) plus final 8 μg/mL polybrene in the same T25 flask followed by incubation overnight at 37 ^ͦ^ C (5% CO_2_) for an additional 24 hours. After 48 hours, we replaced the HEK293T media with 10% FBS DMEM supplemented with puromycin at a final concentration of 2 μg/mL to select for cells which successfully stably integrated our protein overexpression constructs. HEK293T were continued in puromycin for 2 passages until the mock transfected cells died from negative selection in puromycin. We verified successful GFP overexpression in HEK293T cells via a Diaphot Phase Contrast 2 microscope and Olympus DP21 microscopy camera (Nikon, Cat: ELWD 0-3 | Olympus, Cat: DP21). 6-well plates of the newly established AR, SOX2, and GFP-overexpressing cells were plated (5 x 10^6^ cells/well) in 6-well plates and had their protein overexpression verified at 48h via western blot as described previously.

### Nano-BiT assays

We acquired mammalian overexpression constructs encoding the entire full-length proteins of AR, FOXA1, and SOX2 plus either small or large-BiT tags to their N- and C-termini cloned into the pcDNA3.1(+) plasmid backbone from GenScript (See key resources table). To generate the R219-mutations in the FOXA1 plasmids, we performed site-directed mutagenesis using the NEB Q5 kit (New England Biolabs, Cat: E0554S) using primers [FOXA1-R219S-F: GAACTCCATCAGCCACTCGCTGTC, FOXA1-R219S-R: TGCCAGCGCTGCTGG, FOXA1-R219C-F: GAACTCCATCTGCCACTCGCTGT, FOXA1-R219C-R: TGCCAGCGCTGCTGG] according to manufacturer’s protocol. We first these transformed vectors into DH5α *E. coli*, which were then streaked for isolation 1.2% LB agar plates containing 100 μg/mL ampicillin and incubated at 37 ^ͦ^ C for 16 hours. Colonies were picked and grown at 37 ^ͦ^ C for 16 hours in 250mL LB/ampicillin broth for 16 hours. We then isolated and purified our overexpression vector plasmid(s) from the bacteria using a MaxiPrep kit (ThermoFisher, Cat: K0491) eluted in TE buffer and verified concentrations/purity via NanoDrop assay. All plasmids were stored at −20 ^ͦ^ C unless otherwise stated. HEK293T (AR, SOX2, or GFP-overexpressing) cells were plated in clear-bottom poly-d-lysine coated 96-well plates (Corning, Cat: 3603) at a density of 3,000 cells/well in 100μL of 10% FBS DMEM + 2 μg/mL puromycin. Cells were incubated at 37 ^ͦ^ C (5% CO_2_) for 48 hours to attach. We co-transfected in BiT plamids using Turbofectin 8.0 reagent (OriGene, Cat: TF81001) according to manufacturer’s protocol (final 100ng plasmid/well). 6 hours after the initial transfection, the 96-well plates were supplemented with whole 10% FBS DMEM+ 2 μg/mL puromycin. After 12 hours, the media was gently removed using a multichannel aspirator and replaced with 100μL of either 10% FBS or 10% Charcoal-stripped serum (CSS) () phenol-red free RPMI (Gibco, Cat: 11835030) + 1 ug/mL puromycin and returned to the incubator for an additional 24 hours. Next, the wells containing 10% FBS media were treated with 100μL of either DMSO (Sigma-Aldrich, Cat: D2438) or 2X enzalutamide (40μM) to a final concentration of 20μM and incubated for 21 hours. After that time elapsed, the wells containing 10% CSS media were treated with 100μL of either sterile-filtered ethanol (EtOH) (Fisher Chemical, Cat: A962F) or 2X R1881 (2nM) to a final concentration of 1nM and incubated for 3 hours. After 3 hours, all wells had their media carefully removed via multichannel aspirator and replaced by 100μL of 37 ^ͦ^ C Opti-MEM (Gibco, Cat: 11058021). 25uL of Nano-Glo substrate buffer was added per well and the plate luminescence was immediately read using Gen5 3.14 software on the BioTek BioSpa 8 instrument (Agilent, Cat: 11-120-551).

### Chromatin immunoprecipitation and sequencing (ChIP-seq)

CWR-R1 and NCI-H660 cells were plated at a density of 4.1 million cells per IP in 15cm^3^ dishes incubated at 37 ^ͦ^ C (5% CO_2_) for 24 hours. The first grouping of CWR-R1 cells in 10% FBS RPMI (ATCC modification) were treated for 24 hours with either DMSO (Vehicle) or 20μM enzalutamide. In addition, another group of CWR-R1 cells were deprived of androgens for 21 hours in 10% CSS RPMI followed by subsequent treatment for 3 hours with either sterile-filtered ethanol (EtOH - Vehicle) or 1nM R1881. Both CWR-R1 and NCI-H660 cells were fixed in 1% w/v formaldehyde (ThermoFisher, Cat: ICN19404780) for 8 minutes followed by quenching in 125mM glycine (ThermoFisher, Cat: A13816.36) for 5 minutes shaking at room temperature. Following fixation and quenching, we performed chromatin immunoprecipitation, size selection, and purification using the Diagenode iDeal ChIP kit for Transcription Factors (Diagenode, Cat: C01010055) according to manufacturer’s protocol. AR was immunoprecipitated with the kit magnetic beads conjugated to polyclonal rabbit anti-AR (PG-21) antibody (Millipore, Cat: 06-680) or the species-matched rabbit IgG control antibody (Cell Signaling, Cat: 2729). SOX2 was immunoprecipitated with the kit magnetic beads conjugated to polyclonal goat anti-SOX2 antibody (R&D Systems, Cat: AF2018) or the species-matched goat IgG control antibody (R&D Systems, Cat: AB-108-C). FOXA1 was also immunoprecipitated with the kit magnetic beads conjugated to polyclonal rabbit anti-FOXA1 antibody (ThermoFisher, Cat: PA5-27157) or the species-matched rabbit IgG control antibody (Cell Signaling, Cat: 2729). Chromatin was sheared for 40 cycles (30 seconds on/30 seconds off) using a Diagenode Pico sonicator (Diagenode, Cat: B01080010) to a size range of 100-500bp and validated using both 1% agarose DNA gels and an Agilent 2100 bioanalyzer (Agilent, Cat: G2939A). Library preparation and sequencing was performed at the University of Illinois Urbana-Champaign (UIUC) Roy J. Carver Biotechnology Center using the Ovation® Ultralow System V2 DNA-Seq Library Preparation Kit (Tecan, Cat: 402666). The libraries were pooled; quantitated by qPCR and sequenced on one 25B lane for 151 cycles from both ends of the fragments on a NovaSeq X Plus with V1.0 sequencing kits (reads were 150bp in length). Fastq files were generated and demultiplexed with the bcl-convert v4.1.7 Conversion Software (Illumina). Peaks were called using MACS2 and aligned to the human genome assembly GRCh38.p14. ChIP-seq data quality was assessed using FastQC (v0.12.1). IGV (v2.16.2) was used to visualize normalized ChIP-seq read counts at specific genomic loci. ChIP-seq heatmaps were generated in Python using deepTools (v3.5.6) and show normalized read counts at the peak center ±1 kb unless otherwise noted. Overlap of ChIP-seq peaks was assessed using BEDTools (v2.30.0). Peaks were considered overlapping if they shared >= 1bp of overlap.

### ChIP-seq analysis

ChIP-seq peak BED files were analyzed using the web platform for Cistrome-GO^98^ in solo mode to identify functional enrichment of our ChIP-seq peaks using the GRCh38.p14 chromosome build. The custom Cistrome-GO cutoffs were set at <10kb distance from TSS and an adjusted regulatory potential score (adjRPscore) of >= 0.01 to generate lists of candidate target genes for each transcription factor. These lists of nearby genes were then used to generate functional Venn diagrams of shared gene targets. These gene lists were then imported and analyzed through Enrichr^99^ and the significantly enriched GO Biological Processes pathways with *p*-values <= 0.05 were used to generate both Venn diagrams and bar plots for each transcription factor in both the CWR-R1 and NCI-H660 ChIP-seq analyses.

### ChIP-seq Motif Discovery and Enrichment

Peak BED files generated from the ChIP-seq were analyzed using the web-based MEME Suite tools (MEME-ChIP, STREME, and Tomtom v5.5.8)^100–103^. MEME-ChIP was set to discover the 5 most enriched motifs by e-value using the HOCOMOCO (Human + mouse orthologs) DNA motif reference dataset. Both STREME and TomTom analyses were performed at default settings with *p*-value cutoffs of <0.05.

### RNA-sequencing analysis

Previously published publicly available RNA-seq data (de Wet,et. al)^25^ was accessed via NCBI GEO (Accession ID: GSE166184). Differential gene expression matrix files containing TPM values for three replicates of untreated control (Control_1, Control_2, and Control_3) and three replicates of SOX2^KO^ cells (SOX2KO_1, SOX2KO_2, and SOX2KO_3) were used to create heatmaps of differentially expressed genes in Python with myGene^104^ (v3.2.2).

### RT-qPCR

Total RNA was isolated using Trizol reagent (Invitrogen, Cat: 15596026) and purified using the Rneasy Mini Kit (Qiagen, Cat: 74104) following the manufacturer’s protocol. For RT-qPCR, 1μg RNA was reverse-transcribed using the Applied BioSystems High-Capacity cDNA synthesis kit with random primers (Applied Biosystems, Cat: 4368814). Synthesized cDNA was diluted 1:5 in H_2_O and amplified for RT-qPCR (n = 3 replicates) using Fast SYBR green master mix (Applied Biosystems, Cat: 4385617) with 250nM gene-specific primers [*SOX2*-F: GCGGAAAACCAAGACGCTC, *SOX2*-R: TCAGCGCGTAACTGTCC, *ROR1*-F: AGTGCTGAATTAGTGCCTACCT, *ROR1*-R: TCGAGGGTCAGGTAAGAATCTTTG, β-actin-F: CACCATTGGCAATGAGCGGTTC, β-actin-R: AGGTCTTTGCGATGTCCACGT]. β-actin was used to normalize RNA input. Real-time PCR was performed using an Applied Biosystems QuantStudio 3 instrument (Applied Biosystems, Cat: A28231). The mRNA expression levels were calculated using the comparative CT method (ΔΔC_T_).

### Computational modeling of protein interactions

We used the Chai-1 build of AlphaFold-3 to predict the structure of FOXA1 and SOX2 bound to a synthetic promoter (UniProt IDs: P55317 and P48431 respectively)^105,106^. Models were generated without template constraints with 200 diffusion steps and 6 recycles. Confidence in atom placement was calculated as predicted local distance difference test (pLDDT).

### Single-cell RNA-seq Data Acquisition and Preprocessing

Single-cell RNA sequencing (scRNA-seq) data from castration-resistant prostate cancer (CRPC) patient tumors were obtained from the Human Metastatic Prostate (HMP) dataset from Chan et al. 2022 (GSE210358)^70,71^. All processing was done using Seurat (v4.3.0). Low-quality cells were excluded based on total read count and the proportion of mitochondrial genes. The data were normalized using the “LogNormalize” method from Seurat and subsetted to include only tumor cells. Three data files were then utilized: (1) log-normalized gene expression matrix, (2) cell-level scoring data including AR and NE signature scores, and (3) UMAP coordinates with associated metadata. All datasets were processed using R (v4.3.2) and the following libraries: ggplot2 (v3.4.4), ggnewscale (v0.4.9), reshape2 (v1.4.4), ggh4x (v0.2.5), and tidyverse (v2.0.0). Identifiers for each cell were harmonized by renaming columns and coercing types. Expression values for genes of interest (*SOX2*, *FOXA1*, *AR*) were extracted and transposed where necessary for integration with cluster annotations and metadata.

### UMAP Visualization and Gene Expression Overlay for Single-Cell RNA-seq

UMAPs were generated with Seurat (v4.3.0) using the top 2000 features. Individual cells were scored by taking the sum of all genes found from a given signature and then multiplying the sum by the fraction of genes in the signature that had non-zero expression. Cells were annotated according to sample ID, CRPC subtype, AR and NE signature scores. To allow for multi-gene visualization, separate color scales were implemented using the ggnewscale package.

### Quantitative Cluster-Level Expression Analysis and Bubble Plot Construction

To analyze transcriptional programs across tumor clusters, expression data were merged with cluster identity and CRPC subtype metadata. Mean expressions for *FOXA1*, *SOX2*, *AR*, AR signature score, and NE signature score were computed for each cluster-subtype pair. The aggregated values were min-max normalized within each variable to generate a relative scale. Normalized expression data was aggregated and visualized using a bubble plot, where bubble size represented relative expression and color denoted the gene or score.

### Bulk RNA-Seq Signature Scoring and Correlation Analyses

Log_2_-transformed and TPM-normalized bulk RNA-seq data from CRPC-adenocarcinoma samples were obtained from the Beltran et al. 2016 dataset^40^. Signature scores were calculated using a normalized aggregate expression method from gene sets representing luminal activity, basal activity, neuroendocrine identity, and AR signaling, as previously described^36^. Briefly, a relative enrichment score was computed for each sample by summing expression values across genes and rescaling the total to a 0–100 scale. Spearman correlation coefficients were then computed between the signature scores and the expression of each gene within the dataset. Snake plots using these sorted spearman coefficients were generated using ggplot2 and ggrepel, with selected neuroendocrine and AR-axis associated genes labeled. This analysis was used to evaluate the association between *SOX2* expression and basal, luminal, and NE lineage programs in Beltran adenocarcinomas and NEPCs. Pairwise gene-gene correlations between *SOX2* and *ASCL1* were similarly assessed in both Beltran adenocarcinoma and NEPC samples. All correlation analyses were performed using the cor() function from the stats R package.

### Pre-Processing of Public Datasets

Beyond the datasets generated within this work as described in other methods sections above, all RNA-seq data were downloaded from public resources and were pre-processed according to the specifications below. Processing was done for each dataset independently depending on the starting format to ensure that final dataset was in in Transcripts Per Million (TPM) before subsequent analyses.

### GTEx Dataset^37^ (Benign Prostate Tissue)

RNAseq data was downloaded directly from the GTEx web portal already in TPM format, therefore, no additional conversion was required. Duplicate gene names were made unique and all genes with zero expression across all samples were removed.

### TCGA Dataset^38^ (Primary Prostate)

Using the TCGA biolinks (version 2.30.4, project = “TCGA-PRAD”) package in R, raw read counts and associated transcript length information were downloaded for the TCGA Prostate dataset. Raw reads were then subset to include only protein-coding genes. Conversion to TPM was then performed to account for within-sample normalization that was based on overall read depth.

### SU2C/PCF 2019 Dataset^39^ (CRPC)

Data was downloaded directly from cBioPortal in the format of Fragments per kilobase of transcript per million mapped reads (FPMK) and then converted to TPM.

### Adeno (Beltran)^40^ and NEPC (Beltran)^41^

Data was downloaded directly from cBioPortal in the format of Fragments per kilobase of transcript per million mapped reads (FPMK) and then converted to TPM.

### ALAN Analyses

We leveraged the ALAN model and its depicted outputs, as based on our prior study^36^. Per the methodology above, all input data for public datasets were converted into TPM to account for within-sample normalization based on read depth before running ALAN. The ALAN profiles of SOX2, AR, and FOXA1 were extracted from the ALAN outputs and visualized using violin plots (**Figure 2B-F and Figure S2H-L)**.

### Violin Plot Analysis of Time Course Data

Time-resolved bulk RNA-seq data were obtained from the PARCB genetically engineered mouse model of prostate cancer, which recapitulates lineage plasticity and transdifferentiation to small cell neuroendocrine prostate cancer (SCNPC), as described by Chen et al. 2023 (GSE240057)^56^. RSEM data was downloaded and processed to assess longitudinal changes in the expression of key transcription factors involved in lineage reprogramming. Genes with zero counts across all samples were excluded. The data was normalized to transcripts per million (TPM) using transcript lengths derived from GRCh38.p13 annotations and were further log-transformed. To investigate temporal regulation of neuroendocrine drivers, *SOX2* and *ASCL1* expression levels were extracted and visualized across six timepoints: Week 0, Week 4, Week 8, Week 10, Week 12, and Week 18. Data were plotted using ggplot2 with violin plots to capture expression distribution dynamics across replicates per timepoint.

### Statistical Analyses

For cell viability experiments in CWR-R1, 22Rv1, and LASCPC-01, raw absolute luminescence units (ALU) values for technical replicates (n=5) of each treatment group were normalized to their respective untreated controls (-DOX) at each time point and analyzed using a two-tailed paired student’s t-test. Asterisks for significance use the following system, unless otherwise noted: ns = *P* > 0.05, * = *P* ≤ 0.05, ** = *P* ≤ 0.01, *** = *P* ≤ 0.001, **** *P* ≤ 0.0001. For cell viability experiments in NCI-H660, raw absolute luminescence units (ALU) values for technical replicates (n=5) of each treatment group were analyzed using a two-tailed paired student’s t-test. In the Nano-BiT luminescence assays, we subtracted the mean background luminescence values non-transfected wells from each of the technical replicate (n=3) wells and assayed significance between groups using a two-tailed paired student’s t-test. MEME and STREME motif analyses report the estimated statistical significance of a motif as an E-value^100,102,103^. Cistrome-GO functional ChIP-seq peak analyses were performed in solo mode, and lists of candidate target genes were filtered for the genes which fell within 10kb of ChI P-seq peaks and had an adjRPscore of >0.01. Gene ontology pathway enrichment was performed using Enrichr with a *P*-value cutoff of <0.05.

## Data and code availability

Code for this project can be found here: https://github.com/1jphoenix/Phoenix-SOX2-FOXA1

## Acknowledgements

We would like to thank the Prostate Cancer Foundation, and Dr. Howard Soule, PhD and Dr. Andrea K. Miyahira, PhD. We would also really like to thank the Department of Cancer Biology at Loyola University Chicago and the support of the chair, Dr. Nancy Zeleznik-Le, PhD. We would also like to acknowledge the generous support of the Cardinal Bernadin Cancer Center, and its director William Small Jr., MD, FACRO, FACR, FASTRO. We thank Dr. Jiwang Zhang, MD, PhD and Govinda Hancock for generously providing reagents and supplies. We thank Father Peter Breslin, PhD, Dr. Emma Feeney, PhD, and Dr. James Lodolce, PhD, for their work in directing the BIOL 390 lab course and support of the Cardinal Bernadin Cancer Center undergraduate summer training program. We thank Dr. Peter Nelson, MD of UW Fred Hutchinson Cancer Center for providing LNCaP-APIPC and LNCaP-shAR cells. We thank Nick Achille, Dr. Alvaro Hernandez, PhD of the UIUC Roy J. Carver Biotechnology Center, Carlos Martinez and Dr. Mark Maienschein-Cline, PhD of the Research Informatic Core at UIC for their technical support with ChIP-sequencing. We also thank Dr. Mitchell Denning, PhD and Madeline Tara for support with microscopy, as well as Emily K. Mirabella, JD, for her assistance proofreading and editing the manuscript. The results shown here are in whole or in part based on data generated by the TCGA Research Network: https://www.cancer.gov/tcga.

---

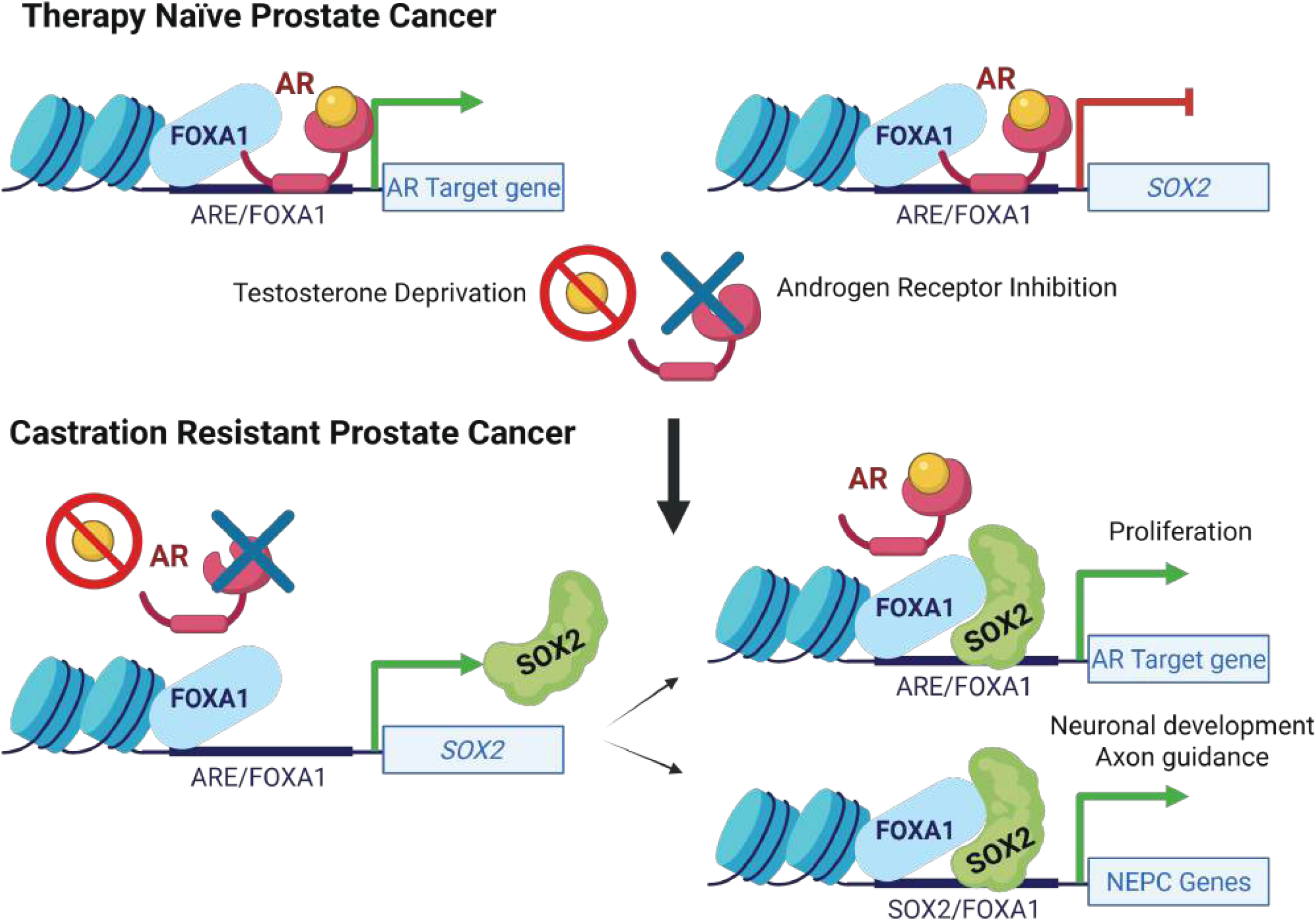

